# Glider compositionality and information capacity in an E/I-balanced neuronal automaton

**DOI:** 10.64898/2026.10.27.761782

**Authors:** Michael T. Belluscio, Sinisa Pajevic, Dietmar Plenz

**Affiliations:** Section on Critical Brain Dynamics, National Institute of Mental Health, Bethesda, MD, USA

## Abstract

In cellular automata, localized activity patterns called gliders propagate and interact to carry and transform information. Whether neuronal networks can harness this computational power remains unclear. Here, we demonstrate that a cortical network model with separate excitatory and inhibitory populations, graded cellular activity and uniform local connectivity spontaneously generates diverse, interacting gliders. An analytical reduction identifies the cellular conditions sustaining propagation of the dominant G-glider. Distinct gliders can assemble into growing structures that emit other glider types. Near excitation–inhibition balance, sparse glider dynamics coexist with high computational capacity, measured by input differentiation and generalization, with local capacity varying across space and time. Strong mutual inhibition between interneurons instead permits metastatic proliferation with low computational capacity. These findings connect cellular automata to neuronal information processing and identify interacting gliders as a candidate mechanism for dynamic plasticity: the reorganization of computation within an unchanged synaptic architecture.

## Introduction

Cellular automata and neuronal networks offer complementary approaches to understanding how complex information processing emerges from interacting units. Neuronal networks describe how synaptic interactions organize collective activity to represent and transform inputs. In cortical networks, excitation and inhibition shape collective dynamics ^1,2^, including oscillations ^3^, traveling waves ^4,5^ and neuronal avalanches ^6,7,8^. Cellular automata reveal how simple interaction rules applied to neighboring cells can generate gliders: localized activity patterns that repeatedly re-form at displaced positions ^9,10,11,12^. Gliders carry information through space, and their collisions can be utilized to implement logical operations, as demonstrated by gates and arithmetic circuits in Conway’s Game of Life ^13,14^. Local information-transfer measurements identify these propagating structures as carriers of information ^15^. In evolved binary cellular automata, propagating particles and their interactions implement collective computation, including density classification ^16^. Their computational appeal lies in this combination of diversity and persistence of their identities, together with their mobility and interactions. Connecting the two approaches requires identifying how neuronal excitation and inhibition generate gliders and determine when they support computation.

Experimental evidence that the cortex can preserve a diversity of activity patterns during propagation comes from the finding of ‘coherence potentials’^7,17^. Recordings in awake monkeys and cortical cultures show that, above a local activity threshold, complex local field potential (LFP) waveforms generated by neuronal populations and their local spiking activity recur at distant sites with little distortion. Their preservation depends on excitatory and inhibitory synaptic transmission ^7,17^. Coherence potentials therefore suggest that glider-like dynamics may be biologically realizable: a pattern can retain its organization as activity passes between neuronal populations. Explaining how cellular interactions sustain this property would connect a computational principle of cellular automata to an observed feature of cortical dynamics.

Prior models establish several parts of this connection. Synfire chains transmit activity reliably through embedded neuronal pathways, including in balanced networks with organized inhibition ^18,19^. Excitable and complexity-based cellular automata support mobile patterns ^20,21,22^. Continuous cellular automata extend cell states from discrete values to a continuous range ^23^, including Lenia, where graded local interactions sustain propagating structures ^24^. Compact propagating gliders in balanced excitation–inhibition networks have been shown to change course when approaching each other^25,26^ or in response to external inputs^27^, providing a substrate for cascading computation. Glider-based logic has also been constructed in excitatory– inhibitory threshold networks through targeted rewiring ^28^.

The computational role of gliders depends on how their propagation and interactions shape neuronal responses to inputs. Reservoir computing provides a framework for relating evolving network activity to input history ^29^, including in cellular automata ^30^, and for assessing whether responses distinguish different inputs while generalizing across related inputs ^31^. This shifts attention from the persistence of individual patterns to the organization of collective activity: how differentiated dynamics remain coordinated across a network, as captured by neural complexity ^32^, and how intrinsic correlations relate to input statistics, as assessed by matching complexity ^33^. For gliders, this organization can change through interactions that annihilate existing patterns or construct new ones. Such composition raises a question of emergence: whether interacting gliders generate collective dynamics with predictive information beyond that available from their constituents separately ^34^. Beyond processing external inputs, spontaneously emerging gliders could initiate and sustain internally organized activity sequences. In the brain, these dynamics could provide a substrate for internally generated thought that is not tied to a particular input. The computational significance of neuronal gliders therefore encompasses both their responses to external signals and their potential to generate structured activity from within, while distinguishing these capacities from uncontrolled proliferation.

Here, we demonstrate that a continuous neuronal automaton with separate excitatory and inhibitory populations and uniform local connectivity supports externally evoked and spontaneous, interacting gliders. We derive analytical conditions for G-glider propagation, characterize the diversity of glider collision outcomes, and show that near excitation–inhibition balance, sparse glider dynamics support high computational capacity that varies across regions and time. We also demonstrate regimes in which glider proliferation is detrimental to computation. This neuronal automaton connects cellular automata and cortical network dynamics through a shared computational resource: self-sustaining patterns of activity that propagate and interact.

## Results

### Local excitation and inhibition sustain network activity

To test whether local neuronal interactions can sustain coherent propagation, we coupled an excitatory grid of 100×100 cells to an inhibitory grid of 50×50 cells (Fig. 1a). Connections within each population were spatially symmetric. Between populations, inhibitory cells provided symmetric local feedback, whereas the chosen excitatory projection recruited inhibition asymmetrically. Periodic boundaries maintained spatial homogeneity. The two grids represent excitatory and inhibitory populations, rather than two anatomical cortical layers.

**Figure 1.**
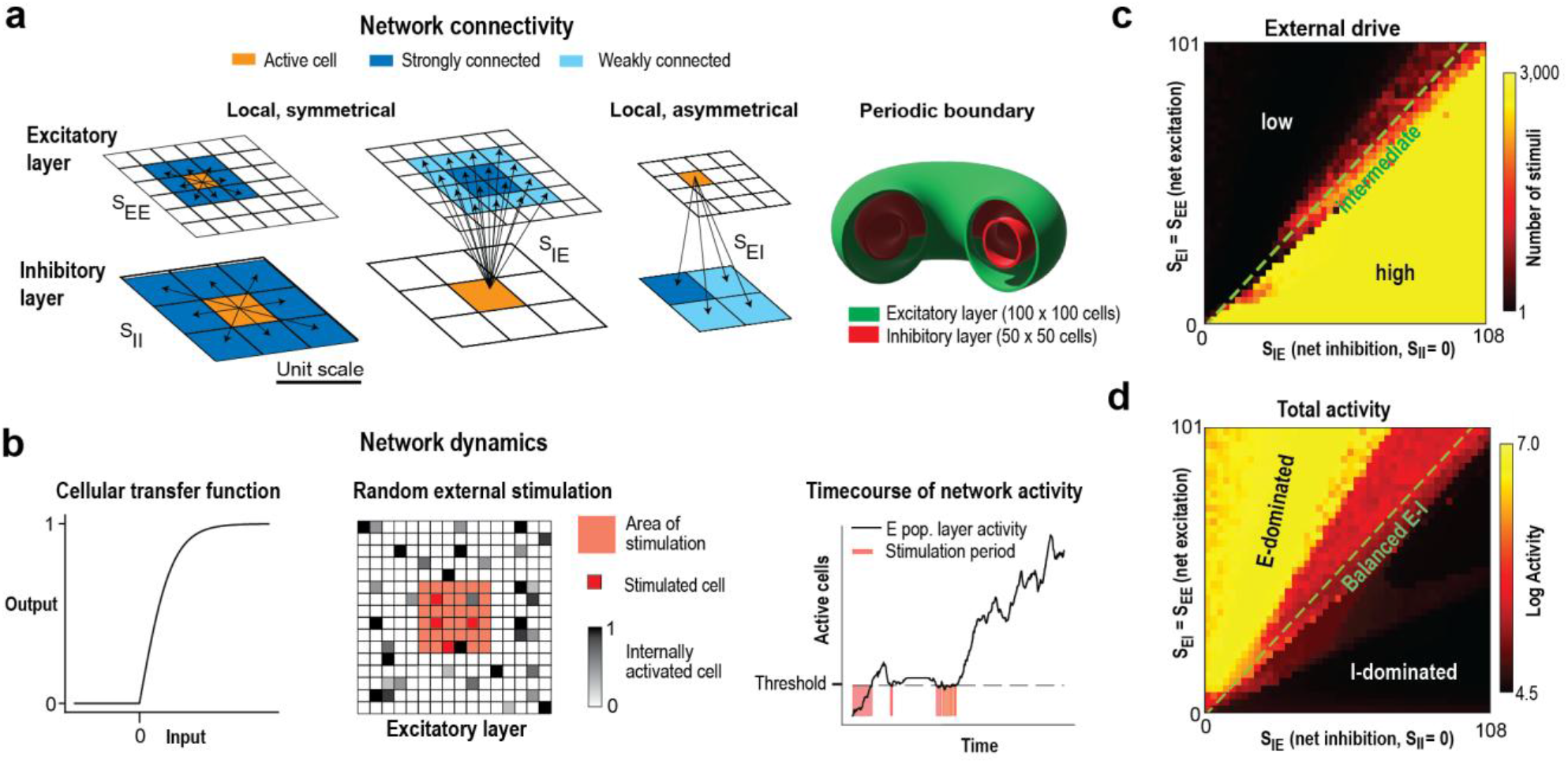
Local excitation and inhibition support distinct activity regimes. (a) Excitatory and inhibitory populations occupy periodic grids with a 4:1 cell ratio. Within-population connectivity is spatially symmetric; the chosen E→I projection is locally asymmetric. Strong interpopulation connections have twice the weight of weak connections. (b) Rectified transfer function, central stimulation region and activity-dependent drive based on the excitatory active-cell count. (c,d) External stimulation count and total activity across S_EE_ = S_EI_ and S_IE_, with S_II_ = 0. The dashed diagonal marks equal coupling. Each condition comprises eight simulations of 3,000 time steps.

Each cell had a continuous activity state between zero and one, updated from its summed input at discrete time steps on a fixed lattice (Fig. 1b). A cell remained refractory for three subsequent time steps after activation. Adaptive stochastic stimulation supplied activity within a central 12×12 excitatory region when the active-cell count was low. The remaining dynamics were deterministic. This protocol permitted persistent activity while avoiding continuous drive in regimes that could sustain their own activity.

We covaried recurrent excitation and excitation of inhibitory cells, S_EE_ = S_EI_, against inhibitory feedback, S_IE_, initially with inhibitory–inhibitory coupling S_II_ = 0. An intermediate regime requiring intermittent drive separated excitation-dominated activity from inhibition-dominated suppression (Fig. 1c,d). We refer to this intermediate region near the equal-coupling line as the balanced regime. This is an operational description of the model’s dynamics and coupling parameters; equality of coupling strengths alone does not establish cancellation of synaptic inputs at individual cells.

### Gliders emerge near excitation–inhibition balance

We next searched for localized activity patterns that recur at displaced positions: gliders. Candidate gliders had to exhibit two successive recurrences with the same period and displacement (Supplementary Fig. 1). We frequently detected a ‘G-glider’ that repeats its spatial configuration every four time steps and moves diagonally at half a cell per time step along each axis (Fig. 2a). Its cycle alternates between a G-shaped configuration of eight active excitatory and five active inhibitory cells and a minimal state of two adjacent active excitatory cells. After two steps the pattern is reflected across its propagation axis; after four steps its orientation is restored.

**Figure 2.**
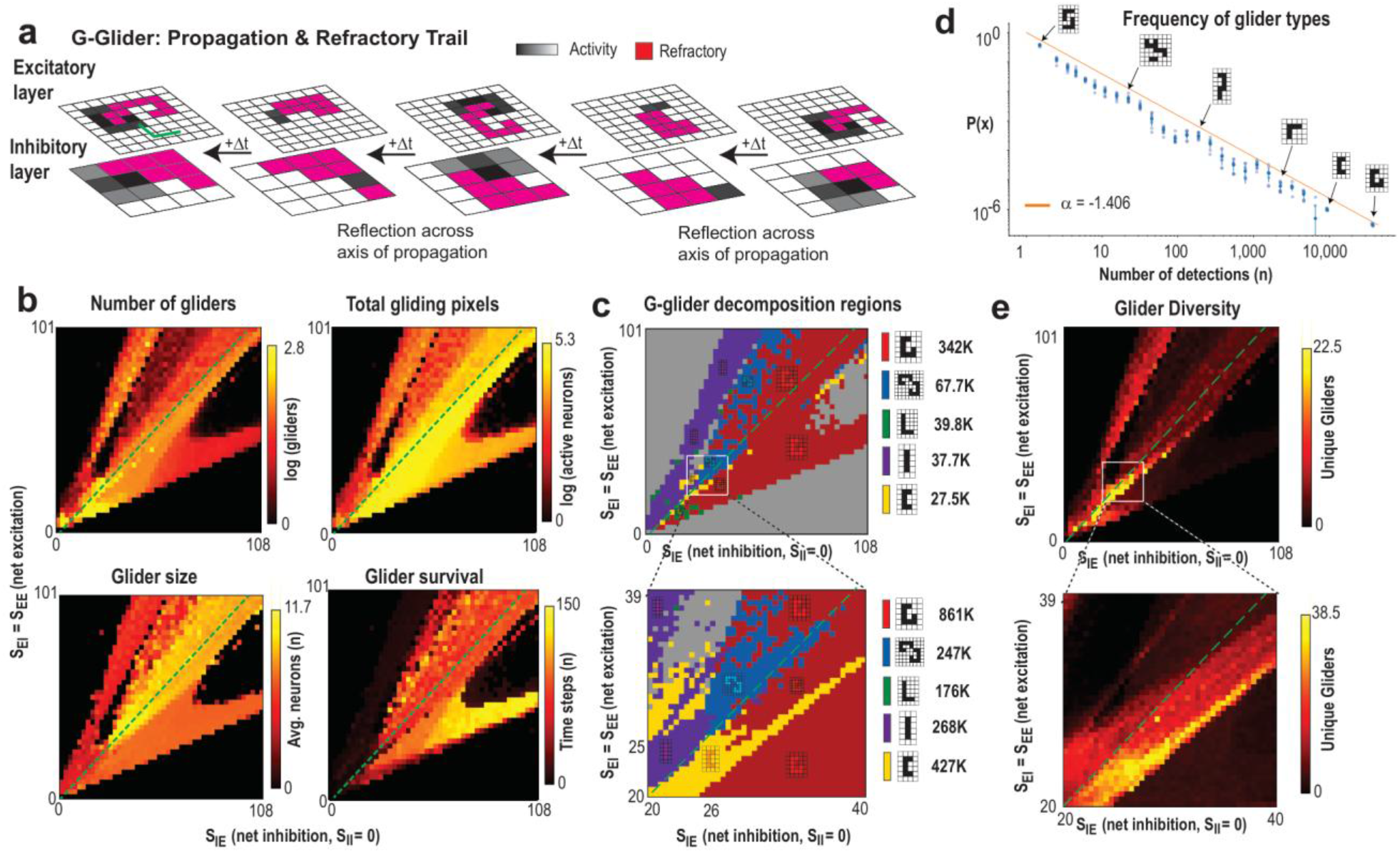
Gliders occupy distinct domains near excitation–inhibition balance. (a) Four-step G-glider cycle, read from right to left, showing active and refractory cells in both populations. (b) Glider count, cumulative gliding activity, mean size, and survival across coupling strengths. (c) Dominance of selected glider patterns and an enlarged view of the low-coupling region; adjacent numbers report detections. (d) Distribution of detection counts across distinct patterns with P(x) representing the relative density of glider types in each detection count bin; the displayed fit has exponent −1.406. (e) Number of distinct patterns detected across parameter space and an enlarged view. S_II_ = 0; eight simulations per parameter point.

Inhibition and refractoriness act together during this cycle. The inhibitory component restricts activation to the two-cell state. Release from that inhibition permits neighboring nonrefractory cells to regenerate the G-shaped configuration of excitatory cells. The surrounding refractory pattern constrains which cells can participate in the next transition and thereby sustains directional propagation. Detection-aligned averages revealed reduced activity ahead of G-glider emergence, beginning seven to eight time steps before detection (Supplementary Fig. 2). This preceding quiescent region persisted under a relaxed matching criterion, although a detection-conditioned average does not establish that the network causally prepares a propagation route.

Glider production concentrated near the balanced region (Fig. 2b). We characterized abundance, size, survival, and cumulative gliding activity. Weak balanced coupling produced many short-lived gliders, whereas stronger coupling produced fewer, longer-lived gliders. This association is consistent with more frequent destructive encounters in dense populations. Different patterns occupied distinct dominance regions, with diversity concentrated near their boundaries (Fig. 2c,e; Supplementary Fig. 3).

**Figure 3.**
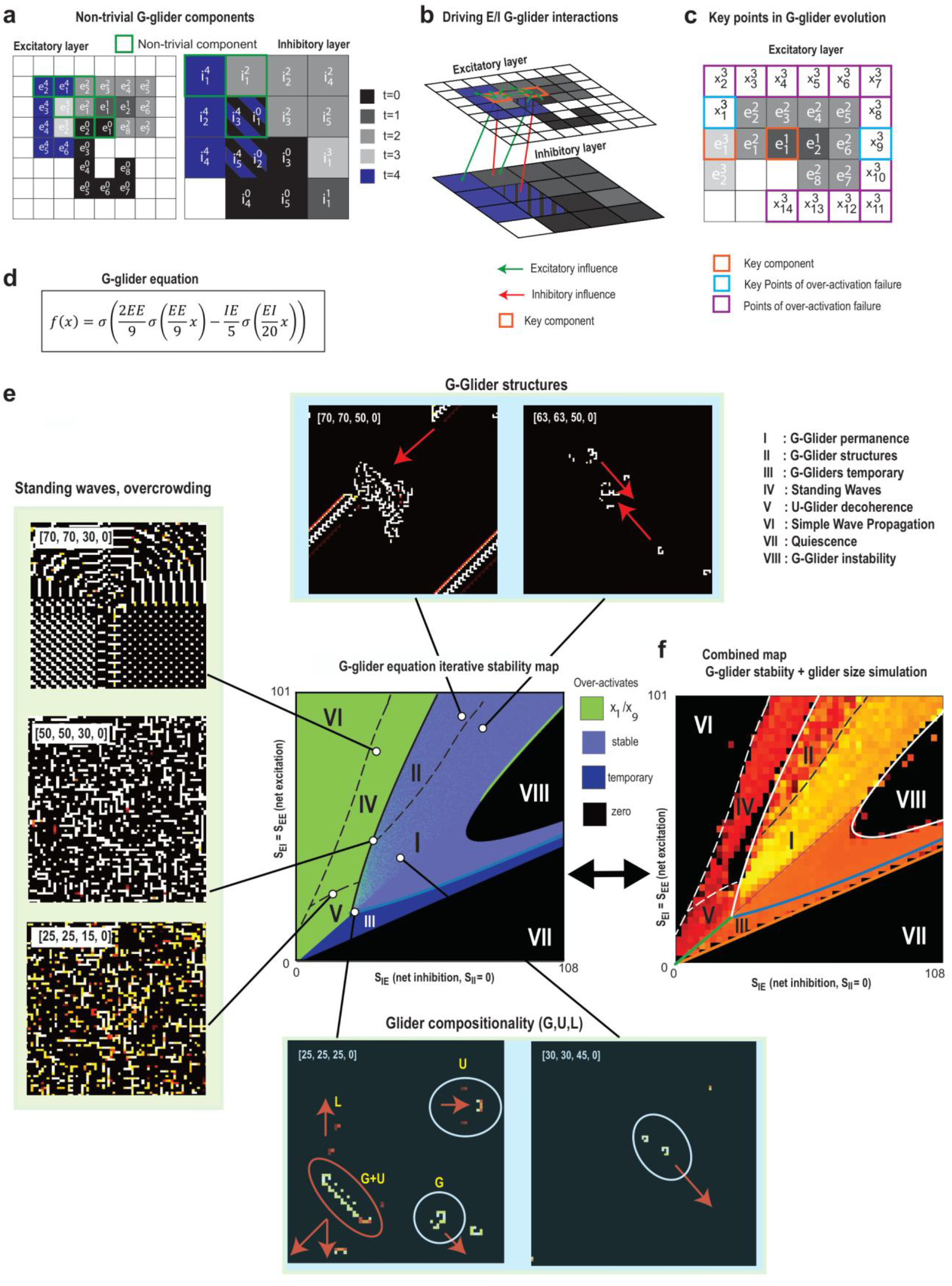
Cellular propagation rules organize glider stability and composition. (a,b) Component cells and their excitatory and inhibitory interactions retained in the reduced G-glider map. (c) Sites at which additional activation can disrupt the assumed spatial pattern. (d) Map for the activity of a propagating excitatory component; one iteration advances half of the four-step cycle. (e) Iterative stability map with representative network snapshots. Light blue denotes stable amplitude dynamics, dark blue temporary/decaying activity, black zero activity and green off-pattern activation. The eight regions distinguish persistent G-gliders (I), growing structures (II), decaying G-gliders (III), standing waves (IV), the U-glider regime (V), simpler waves (VI), quiescence (VII) and G-glider instability (VIII). Upper snapshots show G-glider structures; left snapshots show standing-wave and overcrowded activity; lower snapshots show mixed G/U activity associated with L-glider emission and isolated G-gliders. Arrows indicate the source propagation directions. (f) Reduced-map boundaries overlaid on the simulated glider-size map. Solid boundaries: analytical map. Broken boundaries: domains identified from the full model simulations. Coupling tuples are ordered (S_EE_,S_EI_,S_IE_,S_II_). See Supplementary Note 1 and Supplementary Fig. 5.

We grouped detected gliders by their exact activity configuration. The search catalogued 703 distinct detected configurations, with the most abundant detected 3.4 × 10^5^ times (Fig. 2c,d). A few configurations dominated the detection-frequency distribution, while many occurred rarely. The finite search windows and recurrence intervals limit the sampled repertoire (Supplementary Fig. 1a–e). Varyingthe search-window size from 7 to 15 increased the mean number of neurons in detected patterns in several parameter regions, including the compositionality region (Supplementary Fig. 1f). The catalogue therefore samples only part of the available repertoire.

### Cellular interactions define G-glider propagation

A recurring spatial pattern can persist only if its component cells sustain sufficient activity without recruiting cells that disrupt the pattern. We therefore traced the interactions essential for the G-glider’s progression: one excitatory component in the minimal state and three components in the G-shaped state (Fig. 3a–c).

Their dynamics yield a reduced map for the activity x of the propagating excitatory component:

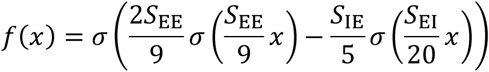

Here σ is the rectified transfer function. One map iteration advances component activity through half of the four-step spatial cycle. A positive fixed point describes a repeated component amplitude; its stability and the absence of disruptive activation impose distinct conditions on propagation. Numerical iteration and checks at sites outside the assumed pattern produced the G-glider propagation map (Fig. 3d,e; Supplementary Note 1). Analytical, numerical and observed boundaries were compared with the simulated glider-size map (Fig. 3f).

The reduced-map analysis corresponded closely to the simulated glider-size map. It distinguished regions of persistent G-gliders, growing G-glider structures, and decaying G-gliders, together with wave regimes and regions lacking sustained propagation (Fig. 3e,f). Amplitude dynamics also exhibited period doubling (Supplementary Fig. 4). These reduced-map dynamics should be distinguished from the stability of the complete spatial pattern and from chaos in the full network.

The full dynamics organized into eight regions (Fig. 3e,f; Supplementary Fig. 5). Persistent G-gliders dominated region I; region II supported growing chains and expanding structures of G-gliders; and G-gliders decayed in region III. Regions IV and VI supported complex standing waves and simpler wave patterns, respectively. U-gliders emerged in region V, where their collisions could generate activity that curtailed further glider propagation. Regions VII and VIII lacked sustained activity under the tested conditions. Boundaries between these regimes supported the mixtures and interactions examined below.

We tested how this organization depended on the cellular rules. Removing refractoriness eliminated gliders in the explored parameter plane. Refractory periods of at least three time steps supported rich glider activity across a broad parameter region within the tested range; shorter periods supported only restricted domains of isolated gliders (Supplementary Fig. 6). Increasing transfer-function gain rescaled the coupling strengths associated with corresponding reduced-map dynamics (Supplementary Fig. 7); changing the transfer-function shape altered their stability and periodicity. The tested linear transfer function did not sustain bounded G-glider activity in the reduced analysis: active solutions amplified without bound (Supplementary Fig. 8). Nonzero inhibitory–inhibitory coupling preserved the broad organization while shifting activity and pattern-specific domains (Supplementary Figs. 9 and 10). An alternative coupling plane provided a complementary view of these dependencies (Supplementary Fig. 11). In this plane, a narrow parameter region showed persistent, metastatic G-glider dominated activity (see below).

### Distinct gliders form growing composite structures

Having identified persistent gliders and their propagation domains, we asked whether different glider types could form larger structures. Most regions supported independently propagating patterns or composites of the same type. Near the boundary between the G- and U-glider regimes, distinct gliders instead formed a connected propagating structure (Fig. 3e, lower snapshots).

The U-glider propagates along a lattice axis (Supplementary Fig. 1b), whereas the G-glider propagates diagonally. In the mixed structure, one U-glider was flanked by one or two G-gliders. As the constituents moved in different directions, the distance between them increased, yet a bridge of activity continued to connect them. The assembly therefore grew as it propagated. Activity at its edge generated L-gliders, leaving a trail of a third pattern type (Fig. 3e, lower-left snapshot). The composite thus combined identifiable propagating components and generated an output distinct from either constituent.

We use compositionality to describe the assembly of recognizable dynamical components into an interacting structure. The growing composite differs from an isolated glider of fixed spatial extent. Its constituent motions, connecting activity and emitted patterns provide an operational basis for assessing composition. This result establishes structural composition in the model, without establishing a general compositional algebra or universal computation.

### Glider collisions generate self-sustaining activity

To determine how interactions reshape ongoing activity, we first tested glider transmission through spatially uniform Poisson noise. A G-glider source launched either isolated gliders or chains of gliders, and transmission was scored by shape-matched arrival at the opposite corner at the expected propagation time (Supplementary Fig. 12). At (S_EE_,S_EI_,S_IE_,S_II_) = (50,50,60,0), chains were more robust than isolated gliders. At (60,60,50,0), the ordering reversed. Transmission therefore depends on both the background coupling regime and the temporal organization of incoming gliders.

We examined 116 collision configurations spanning relative timing, handedness and spatial alignment (Fig. 4a–c). Outcomes included mutual annihilation, survival, redirection, new gliders, composite structures and chain reactions. Some collisions remained transient; others initiated spreading activity that produced a sustained collective state. For each arrangement, the matrices summarize the fraction of sampled coupling settings yielding sustained activity, conditioned on settings where at least one arrangement sustained activity (Fig. 4b). Collision geometry and coupling strengths therefore determined whether a local interaction terminated, transmitted or amplified activity.

**Figure 4.**
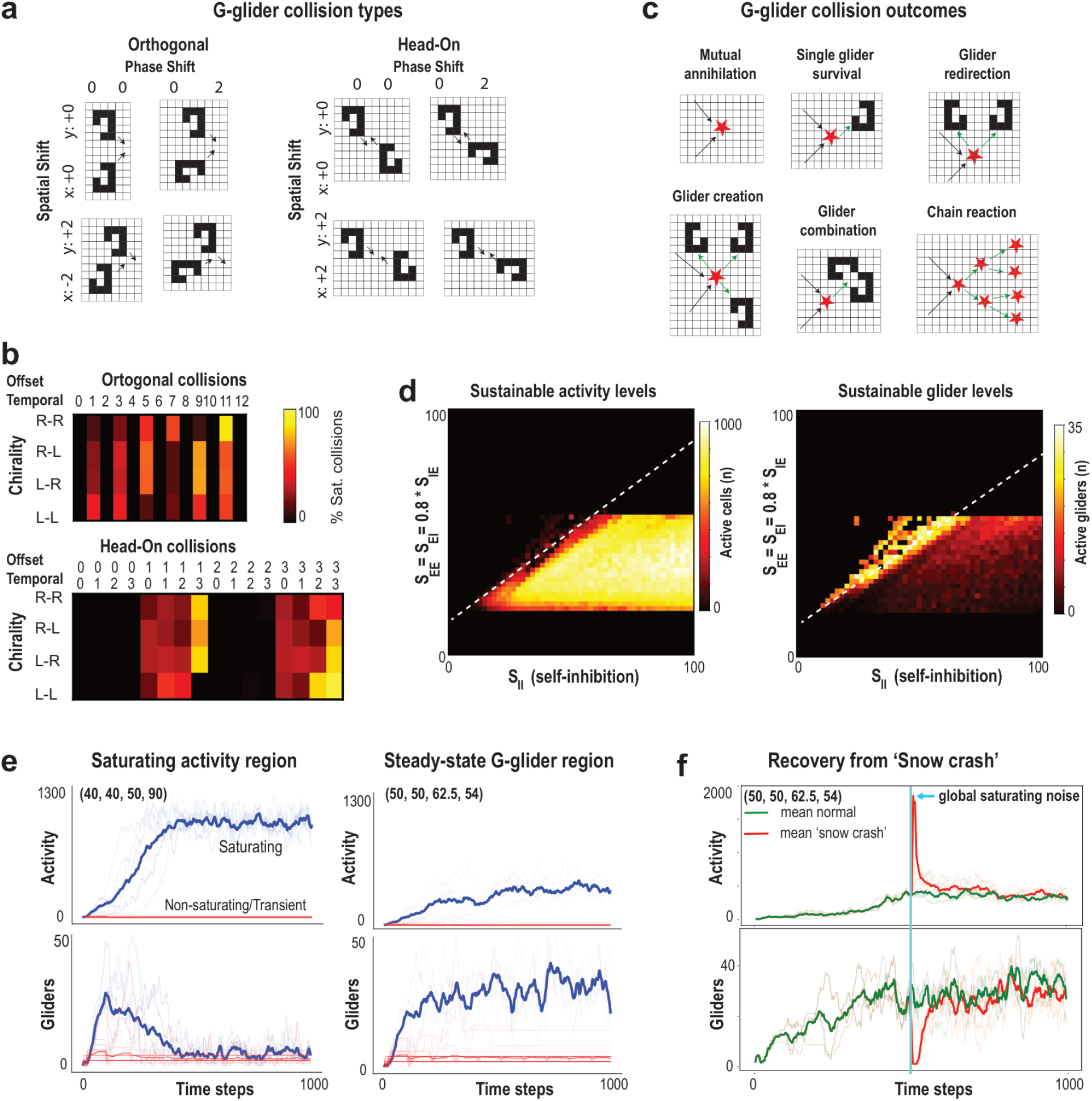
Collision geometry and coupling determine transient or sustained collective activity. (a) Orthogonal and head-on configurations vary phase and spatial offset: 52 orthogonal and 64 head-on arrangements, totaling 116. (b) Configuration matrices display sustained-outcome percentages across the subset of sampled coupling settings where at least one arrangement sustained activity. R and L denote handedness; orthogonal temporal offsets are 0–12, and head-on temporal and spatial offsets are 0–3. These are parameter-scan summaries, not probabilities for randomly sampled collisions. (c) Representative annihilation, survival, redirection, glider creation, combination and chain-reaction outcomes. (d) Sustained activity and G-glider counts across S_II_ with S_EE_ = S_EI_ = 0.8 S_IE_. (e) Activity and glider-count trajectories contrast spreading activity with transient gliders at (40,40,50,90) and persistent gliders at (50,50,62.5,54). (f) Recovery after a brief global perturbation at the latter setting; thin traces show retained arrangements and emphasized curves their means. Tuple order is (S_EE_,S_EI_,S_IE_,S_II_). Network-size dependence is shown only in Supplementary Fig. 13.

Increasing inhibitory–inhibitory coupling permitted collision-induced sustained states over a restricted region of parameter space (Fig. 4d,e). At stronger coupling, gliders appeared transiently during spreading activation and then disappeared. Near the boundary of sustained activity, both population activity and glider counts persisted. Collisions could therefore amplify localized propagation into a collective state, but persistent activity did not necessarily preserve gliders. Neither outcome alone establishes computational capacity.

Sustained activity and glider counts increased with network size for excitatory grids of 52×52 cells and larger; the tested 24×24 grid did not sustain comparable dynamics and is not plotted (Supplementary Fig. 13). A brief global perturbation of noise (‘snow crash’) disrupted the glider population, after which activity and glider counts recovered toward their preceding levels (Fig. 4f).

### Sparse glider dynamics preserve high, time-varying local capacity

The collision results make persistence an insufficient criterion for assessing the computational role of gliders. We therefore asked whether a regime could sustain interacting gliders while retaining differentiated responses to inputs and tolerance to input variation. We used kernel rank and generalization rank to characterize responses to distinct inputs and noisy versions of a common input, respectively ^31^. Neither measure prescribes a task-trained readout.

Kernel rank was computed from responses to 100 distinct sequences. Generalization rank was computed from a reference sequence and 99 noisy variants; we report 100 minus that rank as the generalization measure, so larger values indicate less response variation. Combining kernel rank and the generalization measure by their geometric mean yielded Φ (Fig. 5a,b). High generalization alone can result from quiescence; combining the measures penalizes states that fail to distinguish inputs.

**Figure 5.**
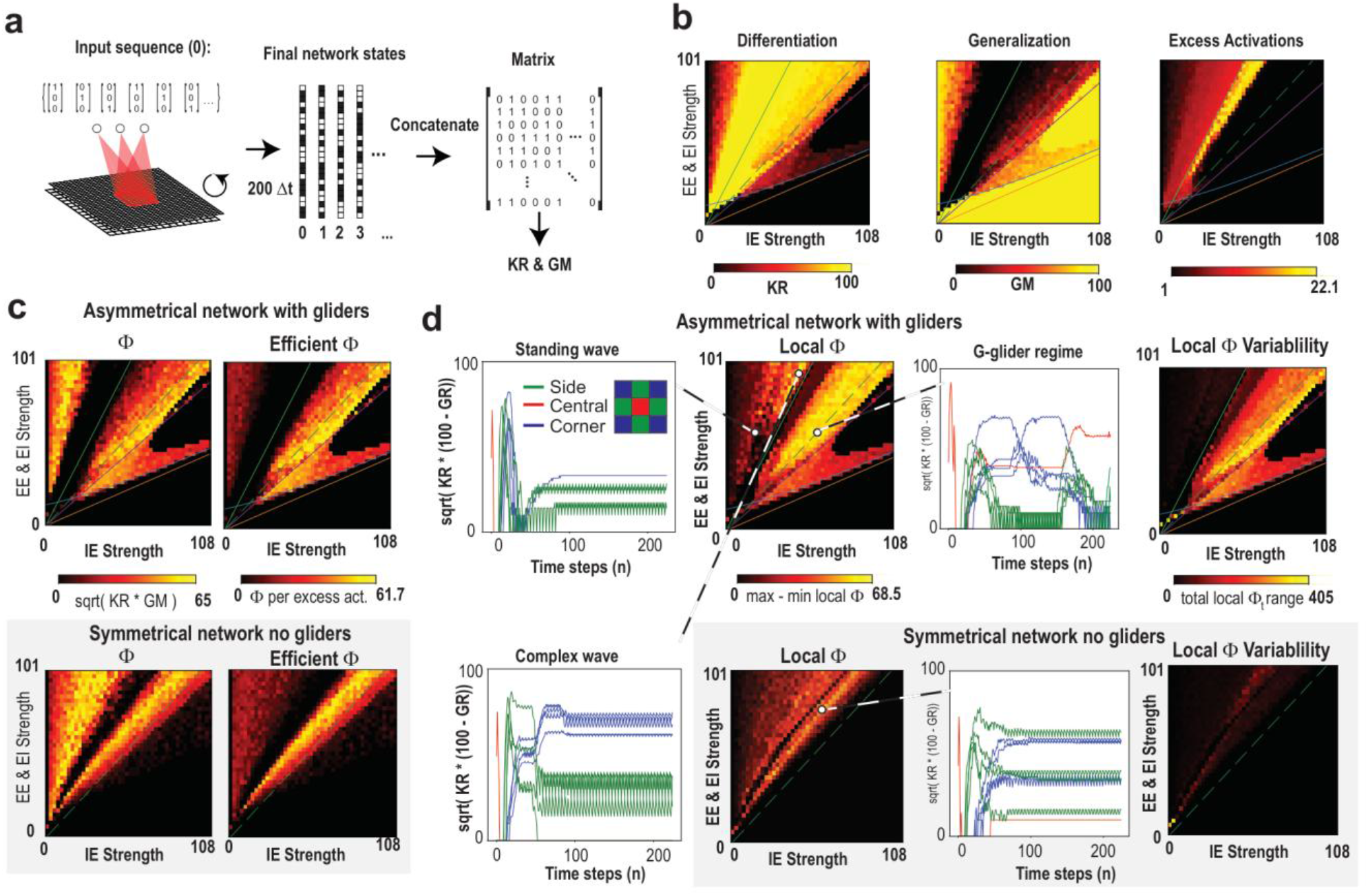
Sparse glider dynamics preserve high, time-varying local capacity. (a) Input sequences and response matrices used to measure kernel and generalization ranks. (b) Kernel rank, the generalization measure and relative activation across coupling strengths. (c) Global Φ and activity-normalized Φ for the asymmetric 4:1 model and symmetric 1:1 control. Φ is the geometric mean of kernel rank and 100 minus generalization rank. (d) Spatial differences and temporal variability of local Φ, with representative regional time series. Local Φ is computed from paired ranks within each region. The spatial reducer takes the maximum over time of the difference between the largest and smallest noncentral regional Φ; the temporal reducer sums each region’s maximum-minus-minimum Φ over time, including the center. Both use samples from time index 100 onward. Red denotes the central subgrid, green side subgrids and blue corner subgrids. The glider example retains changing local capacity after the other displayed examples approach stationary arrangements. The local calculation uses nine 33×33 subgrids within the 100×100 excitatory grid. Inhibitory grids contain 50×50 cells in the 4:1 model and 100×100 cells in the 1:1 control. Eight independent simulations per condition are reported for the maps.

Both wave and G-glider regimes supported high global Φ (Fig. 5c). Their activity costs differed: waves recruited much of the network, whereas glider regimes maintained spatially sparse activity. Dividing Φ by excess activation expresses capacity relative to the model’s activity cost as efficient Φ. High Φ in the glider regime establishes that propagating patterns can coexist with input differentiation and generalization. Gliders therefore need not destroy these capacities.

We next examined when and where this capacity was available. Local Φ was computed separately in nine subgrids of 33×33 cells (Fig. 5d). In the glider example, local capacity continued to vary across regions and time after the initial response. The standing-wave, complex-wave and symmetric-control examples approached more stationary local arrangements. These time series show capacity available locally over finite intervals within the sampled response, rather than a fixed spatial allocation. The spatial summary measures the maximum interregional difference in Φ, and the temporal summary sums the within-region ranges from time index 100 onward.

The symmetric 1:1 control also supported high global Φ despite lacking verified isolated gliders in the reported exploration (Supplementary Fig. 14). Its local capacity reached a lower peak Φ and varied less over time than in the displayed asymmetric glider example (Fig. 5d). A symmetric 9:1 model generated gliders (Supplementary Fig. 15), showing that the particular asymmetry of the 4:1 model is not necessary for all glider emergence.

At the recovery setting, (S_EE_,S_EI_,S_IE_,S_II_) = (50,50,62.5,54), with positive inhibitory–inhibitory coupling, the rank protocol used for Fig. 5 yielded ∼90% smaller Φ. Thus, persistent glider activity can recover in the metastatic regime despite low computational capacity.

Our results thus identify a regime in which sparse glider dynamics coexist with high, changing local capacity, in contrast to the low-Φ metastatic regime. The 1:1 control changes population ratio, inhibitory population size and connectivity geometry together, so it does not isolate a causal contribution of gliders.

## Discussion

We demonstrated that a cortical network model supports a diverse, compositional repertoire of gliders, extending the computational power of gliders in cellular automata to neuronal networks governed by excitation and inhibition. Here, a glider is a cluster of neighboring active cells whose activity pattern re-forms at successive locations as new cells are recruited. An analytical model of the dominant G-glider provides a mechanistic basis for the state-space organization identified in simulations, linking cellular interactions to the boundaries between sustained propagation, decay and recruitment of additional activity.

The model brings together four properties: a power-law distribution of glider occurrence frequencies (Fig. 2); an analytical account of the simulated propagation regimes (Fig. 3); diverse interaction outcomes across 116 collision configurations (Fig. 4); and compositionality, whereby distinct gliders assemble into growing structures that emit gliders of another type (Fig. 3).

These properties acquire functional significance when glider interactions preserve differentiated responses to inputs while allowing generalization across related inputs (Fig. 5). Near excitation–inhibition balance, sparse glider dynamics supported high computational capacity, Φ, with individual regions retaining these capabilities over finite intervals. Wave regimes also supported high computational capacity but recruited more widespread activity, while their local capacity approached more stationary spatial distributions. Glider dynamics therefore combined sparse recruitment with a continually changing distribution of computational capacity. Glider dynamics were not uniformly beneficial for computation: strong mutual inhibition between interneurons permitted metastatic proliferation with low computational capacity.

Together, these findings identify glider dynamics as a potential mechanism for flexible information processing in neuronal networks.

### Architecture and cellular constraints

The continuous neuronal automaton connects cellular-automaton computation to cortical circuitry by combining separate excitatory and inhibitory populations, graded cellular activity and refractoriness with uniform local connectivity. This construction allows gliders and their interactions to emerge from shared cellular rules without embedding dedicated propagation pathways or wiring specific computational operations. The resulting repertoire therefore reveals what local neuronal interactions can generate before introducing the connectivity heterogeneity, synaptic plasticity and external structure of cortical networks.

Recurrent excitation remains spatially symmetric; the local asymmetry enters through the recruitment of inhibition by excitatory cells and thereby shapes recurrent inhibitory feedback. Glider propagation therefore emerges without a learned or selectively wired excitatory pathway, in contrast to the targeted rewiring used for glider-based logic ^28^ and the embedded pathways supporting synfire chains ^18^.

Local asymmetry is not essential for glider emergence: the symmetric 9:1 model also generated gliders near excitation–inhibition balance, where it supported high computational capacity (Supplementary Fig. 15). The coexistence of glider activity and input-dependent computational capacity therefore extends beyond the specific connectivity geometry and population ratio of the 4:1 model.

The rectified, saturating transfer function limits recurrent amplification; the tested linear function instead permits decay or unbounded amplification, without robustly selecting a finite active amplitude (Supplementary Fig. 8). Amplitude regulation and refractory timing reflect fundamental constraints on single-neuron firing: finite firing rates and absolute and relative refractoriness limit how strongly and how rapidly neurons respond. In the model, the G-glider completes its spatial cycle in four time steps, and a three-step absolute refractory period prevents recently active cells from being reactivated during the intervening phases, protecting reconstruction of the advancing pattern (Fig. 2a). Consistent with this mechanism, three steps marked the onset of a broad regime of rich glider activity, whereas shorter refractory periods supported isolated gliders only in restricted parameter regions (Supplementary Fig. 6).

### Glider diversity and analytical insight

Within these cellular constraints, the diversity of glider configurations provides a potential dimension for encoding information through the identity of the propagating pattern. This connects the model to cortical coherence potentials, whose diverse local field potential waveforms reflect distinct temporal patterns of local neuronal firing that recur at distant sites ^7,17^. The power-law distribution of glider occurrence frequencies describes a repertoire of a few common and many rare configurations (Fig. 2d), providing a concrete model of such pattern diversity.

A central advance is the analytical reduction of the dominant G-glider, which closely accounts for its propagation boundaries in the simulated state-space map at S_II_ = 0 (Fig. 3). This agreement shows how a small set of cellular interactions governs the persistence, decay and spatial destabilization of a propagating structure, providing a tractable framework for predicting how these boundaries change with synaptic coupling, neuronal gain and transfer-function shape. The reduction also offers a foundation for studying perturbation sensitivity, identifying other self-consistent glider configurations and developing a statistical theory of interacting glider populations in E/I-balanced neuronal networks.

Whereas the glider equation for Asymptotic Lenia characterizes shape-preserving translation ^24^, our reduction resolves the successive cellular interactions that reconstruct the G-glider during propagation. By linking amplitude dynamics and recruitment outside the pattern to synaptic coupling, it provides a mechanistic account of the propagation boundaries observed in simulations at S_II_ = 0 (Fig. 3).

### Constructive interactions

Particle interactions, including annihilation and transformation, implement collective computation in evolved binary cellular automata ^16^. A diverse glider repertoire provides distinct dynamical components that can combine while retaining their identities, allowing interactions to construct new propagating structures. We demonstrated this form of compositionality in an E/I-balanced neuronal network, where G- and U-gliders form growing connected assemblies that emit L-gliders (Fig. 3).

The collision repertoire further shows that encounters can create new gliders and composite structures or initiate sustained activity, with outcomes determined by collision geometry and coupling strengths (Fig. 4).

Whether these assemblies carry predictive information beyond that available from their constituents separately can be tested using information-decomposition approaches to causal emergence ^34^.

Previous neuronal-network studies demonstrated a form of ballistic computation in which close encounters between compact propagating patterns redirected their trajectories, allowing logical operations based on signal presence or absence to cascade through the system ^25^. Directional alignment quantified collective propagation, while improved stimulus processing was assessed separately through population decoding ^27^. These analyses established signal routing and stimulus representation but left open how interactions could transform pattern identity and construct new dynamical components. Our model demonstrates this constructive dimension: encounters generate new gliders and composite structures, while distinct glider types assemble into growing configurations that emit a third type (Figs. 3, 4).

Glider guns provide recurring signals for logic constructions in cellular automata ^35,36,37^. We also identified a glider gun, a recurrent population pattern that repeatedly emits propagating gliders (Supplementary Fig. 12). Such a structure provides a potential network mechanism for generating recurring spontaneous activity beyond the rhythmic activity of single, ‘pacemaker’ neurons. We note that glider absorbers and constructions aimed at universal computation lie beyond the scope of this manuscript.

### Detection and robustness

Our detection filter identifies activity patterns that recur at displaced locations within defined spatial windows and recurrence intervals (Supplementary Fig. 1a–e). Increasing the search-window size from 7 to 15 increased the mean number of neurons in detected patterns in several parameter regions, including the region supporting glider composition (Supplementary Fig. 1f). This dependence is consistent with larger assemblies extending beyond the main catalogue’s detection scale.

Glider transmission declined gradually under the tested external noise, with robustness depending on synaptic coupling and the temporal organization of incoming patterns (Supplementary Fig. 12). The lattice specifies which neurons interact; it does not require their physical positions to form a regular grid. Permuting neuronal positions while preserving synaptic partners, weights and cell types leaves the model dynamics unchanged under corresponding inputs, even when the spatial arrangement appears disordered.

### Computational capacity and dynamic plasticity

Computational capacity requires a network to distinguish different inputs while generalizing across variations of the same input. Following the kernel-rank and generalization-rank framework of Legenstein and Maass ^31^, we combined these complementary properties in Φ (Fig. 5). This response-based measure assesses the computational potential of the evolving network state without assigning logical operations to individual collisions. This response-based capacity is distinct from the irreducible cause–effect structure assessed by integrated information theory ^38^.

No synaptic weight changed in our model, yet computational capacity shifted across the network. We propose two complementary layers of brain computation: synaptic plasticity, which changes the connections between neurons, and dynamic plasticity, which reorganizes information processing within an unchanged synaptic architecture. Interacting gliders provide a candidate mechanism for this dynamic plasticity, forming transient computational structures through the propagation and interaction of neuronal activity. This dynamic layer also has a failure mode: metastatic glider proliferation sustains activity but supports little computational capacity, separating the persistence of neuronal activity from its usefulness for information processing (Fig. 5).

To summarize, we demonstrated that diverse gliders emerge, interact and form composite structures in neuronal networks, providing a mechanistic link between cellular-automaton computation and neuronal information processing. These findings identify a candidate mechanism for dynamic plasticity, through which information processing can reorganize within a fixed synaptic architecture. We propose that this dynamical level complements synaptic and structural plasticity in supporting cognition, and that its disruption may contribute to cognitive dysfunction.

## Methods

### Network architecture and connectivity

The excitatory population comprised 100×100 cells and the inhibitory population 50×50 cells, with periodic boundaries and a 4:1 cell ratio (Fig. 1a). The ratio captures the predominance of excitatory neurons in cortex; it is a model choice rather than a fixed anatomical ratio ^39^. Within each population, the implemented Moore-neighborhood sum includes the eight nearest neighbors and the cell itself ^40^. The coupling strengths were S_EE_ for E→E, S_EI_ for E→I, S_IE_ for I→E, and S_II_ for I→I.

Each excitatory cell projected strongly to one spatially coincident inhibitory cell and weakly to three nearby inhibitory cells. Each inhibitory cell projected strongly to four coincident excitatory cells and weakly to twelve surrounding excitatory cells. Strong connections had twice the weight of weak connections. This chosen geometry produces local E→I asymmetry while maintaining spatial homogeneity. A 4:1 population ratio alone does not impose this geometry.

### Cellular update

Incoming signals to the excitatory and inhibitory populations were computed as follows:

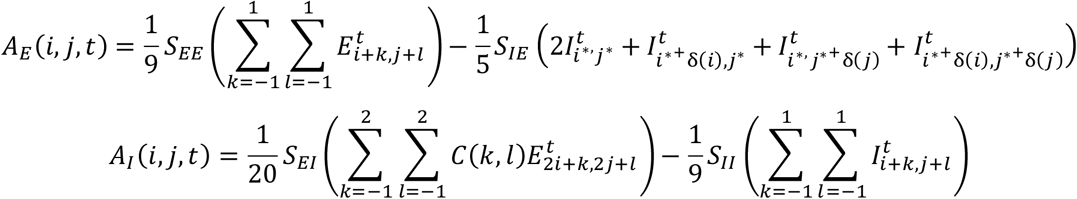

E and I denote real-valued cellular activity states between zero and one, rather than binary activation variables. The normalization factors scale the incoming sums by their total connection weights. Indices between the two grids were related by

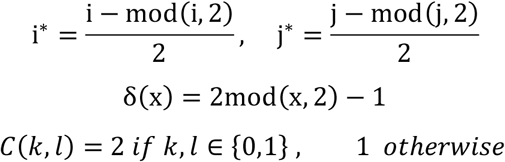

The next state depended on the incoming signal and recent activity:

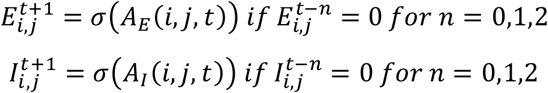

The transfer function was

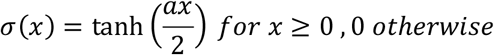

with default a = 1. Refractory cells and cells whose net input fails the activation condition have zero output. A cell with activity greater than zero remains refractory for three subsequent steps. The within-population sums include the center term; refractoriness suppresses immediate self-reactivation. Excitable automata provide a precedent for combining activation and refractory dynamics ^41^.

### Stimulation and parameter exploration

Stimulation targeted one available cell in a central 12×12 excitatory region, with amplitudes sampled uniformly from (0,1]. The production sweeps triggered stimulation after the network update when fewer than 60 excitatory cells were active. Figure 1b uses a fixed color to indicate the stimulation location, not a fixed amplitude.

The principal parameter plane covaried S_EE_ = S_EI_ against S_IE_ with S_II_ = 0. Inhibitory–inhibitory connectivity varies among cortical cell types ^42,43^; its effect was examined separately rather than treated as absent from cortex. The supplied production command specifies 41 equally spaced excitatory strengths from 1 to 101 and 46 inhibitory strengths from 0 to 108: increments of 2.5 and 2.4, respectively, giving 1,886 parameter combinations. Each point comprised eight simulations of 3,000 time steps with independent stochastic stimulation.

The explored nonzero S_II_ values were 2, 5, 10, 20, 30, 40, 50, 80, and 100. Supplementary Figure 10 presents selected pattern-specific maps at 5, 10, 20, 40, 80, and 100. The alternative plane tied S_EE_ to S_II_ and S_EI_ to S_IE_ (Supplementary Fig. 11). The 1:1 control used 100×100 cells in each population. The 9:1 simulations used a 99×99 excitatory grid and a 33×33 inhibitory grid.

Activity was quantified by summing active-cell instances:

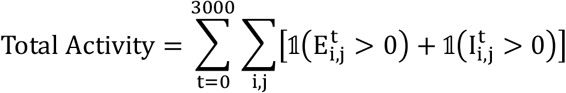

Values were averaged across simulations. Stimulation was counted as the number of external drive events. The supplied activity reducer applies log10(A + 1) to summed activity A before aggregation.

### Glider detection and statistics

We defined an isolated glider as a localized pattern recurring with a fixed displacement after a period τ. The detector binarizes excitatory activity at values greater than zero for pattern matching. This detection step does not change the continuous activity states used in the network dynamics and computational-capacity calculations. It scans rectangular windows of 3–10 cells in each dimension, requiring at least four active cells, a silent one-cell border and an active-cell density of at least 0.2. Candidate translations recur after 1–4 time steps and again after the same interval and displacement. Each displacement component is bounded by τ cells and zero displacement is excluded. The matching tolerance is 0.99; periodic tiling handles boundary crossings (Supplementary Fig. 1). These are detection constraints, not a proof of a general propagation-speed bound.

The detector retains the smallest enclosing window, links repeated detections at the expected translated position, and removes duplicate detections at multiples of a shorter period. Records are retained when linked duration exceeds 2τ. A separate filter rejects searched time points where the entire binary grid recurs at tested period multiples. The supplied validation suite detected a translating localized pattern and a network G-glider, while rejecting its standing-wave, wavefront and stationary-pattern examples. Recurrent fragments of mature standing waves were excluded from the production catalogue; propagating precursors before a standing wave fills the grid can still meet the glider criterion. A separate comparison varied maximum search-window sizes from 7 to 15 to map differences in mean detected pattern size, measured as the average number of active cells across all detected patterns(Supplementary Fig. 1f).

Glider size is the number of active cells in a detected pattern; the displayed size statistic is the mean over detections. Survival is the linked duration in time steps. The supplied cumulative-gliding reducer sums size × duration over detected records and applies log10(total + 1). The mean-size reducer returns zero when fewer than five records are available.

### Noise and collision analyses

Noise transmission was tested by launching gliders from one corner and detecting their shapes at the opposite corner at the expected arrival time. The reported sampling comprised 100 independent single-glider trials and one chain of 100 gliders. Gliders within the chain are not independent trial replicates. For a chain, success is the number of arriving G-glider shapes divided by the number launched; it is not the fraction of chains with at least one arrival. Noise activation counts are sampled from a Poisson distribution with mean λ per network time step; locations are sampled from the permitted grid area and selected cells receive unit activity. At (50,50,60,0), the displayed fits are y = −0.659x + 0.945, r = −0.975 for isolated gliders and y = −0.439x + 1.018, r = −0.918 for chains. At (60,60,50,0), the corresponding fits are y = −0.676x + 0.936, r = −0.959 and y = −0.507x + 0.405, r = −0.589 (Supplementary Fig. 12). The fit intercepts are not observed success probabilities.

The collision catalogue comprises 52 orthogonal arrangements (four handedness combinations and thirteen temporal offsets) and 64 head-on arrangements (four handedness combinations, four temporal offsets and four spatial offsets), totaling 116 (Fig. 4a,b). The returned off-balance plotter samples S_II_ at 51 values from 0 to 100 and S_IE_ at 51 values from 0 to 125, with S_EE_ = S_EI_ = 0.8 S_IE_. It smooths activity over four steps and classifies an arrangement as sustained when the mean absolute difference between initial activity and the final three smoothed samples is at least 25. The configuration-table denominator counts sampled settings where at least one arrangement meets this criterion; the resulting fractions are conditional on this subset. Missing files and empty selections require explicit accounting.

The size-analysis plotter uses (S_EE_,S_EI_,S_IE_,S_II_) = (45,50,50,45), excitatory side lengths 24, 52, 100, 200 and 300, and four-step smoothing. The plotted traces show 52, 100, 200 and 300; the 24×24 condition became quiescent and is omitted (Supplementary Fig. 13). The recovery plotter identifies the setting (50,50,62.5,54) (Fig. 4f). In a 1,000-step run, the perturbation is applied at loop indices 501–510: each quiescent, nonrefractory excitatory cell receives unit activation with probability 0.03 per step. Thin traces show retained collision arrangements and emphasized curves their means.

### Computational measures

For kernel rank, 100 input sequences are sampled uniformly from [0,1]^3^, each lasting 25 time steps. Three input channels project into the central regions of both populations through Gaussian weights of mean zero and standard deviation one. The same projection matrix is reused across the kernel and generalization sequences within a run. Generalization uses the repeated reference sequence [(1,0,0), (0,1,0), (0,0,1)] and 99 variants with Gaussian noise of standard deviation 0.05. Analog excitatory states form the response matrices. The supplied code computes their ranks using NumPy’s default matrix-rank tolerance through the final sample at index 224, after 200 post-input updates. The generalization measure is 100 minus generalization rank.

The weighted input is clipped to zero at values at or below 2, reduced by 2 between 2 and 3, and capped at 1 above 3. Refractory cells cannot receive this drive. These thresholds act on the weighted input itself; they are not a separately estimated input standard deviation.

Global Φ is the square root of kernel rank times the generalization measure. In the activity-normalized reducer, Φ is divided by the geometric mean of the two post-input/input activity ratios, each floored at one. The returned rank driver computes local Φ in nine 33×33 subgrids covering the upper-left 99×99 cells of a 100×100 excitatory grid, storing the central subgrid first. The corrected spatial reducer first computes Φ from paired kernel and generalization ranks in each subgrid. At each sample from time index 100 onward, it finds the maximum-minus-minimum Φ across the eight noncentral subgrids, then takes the maximum across those samples. The temporal reducer sums each subgrid’s Φ range over the same temporal window and includes all nine subgrids. These summaries quantify spatial heterogeneity and temporal variation, respectively.

### Implementation

Simulations were implemented in Python 3 with NumPy. Both populations were initialized at zero activity; time steps are arbitrary units without biological time calibration. The returned package includes the network, detector, reduced-map implementation and original research scripts. Earlier bounded checks covered selected detector examples, analytical boundaries and network updates; they do not validate the changed release or reproduce the production figures.

## Data availability

Source data for the main and supplementary figures will accompany the paper.

## Code availability

The code package contains the neuronal automaton, detector, reduced-map analysis and original research scripts.

## Acknowledgements

We thank Keshav Srinivasan for early discussions on this project as well as other members of the Plenz lab for lively discussions on gliders and their potential benefits. This research was supported by the Division of the Intramural Research Program (DIRP) of the National Institute of Mental Health (NIMH), USA, ZIAMH002797. This research utilized the supercomputing resources of the National Institutes of Health (NIH, USA; Biowulf, http://hpc.nih.gov). This manuscript was assembled and edited with assistance from Claude Code (Opus 5.5; Anthropic) and GPT-6-Astra (OpenAI). The authors take full responsibility for the scientific content and final text.

The contributions of the authors were made as part of their official duties as NIH federal employees, are in compliance with agency policy requirements, and are considered Works of the United States Government.

However, the findings and conclusions presented in this paper are those of the author(s) and do not necessarily reflect the views of the NIH or the U.S. Department of Health and Human Services.

## Author contributions

DP conceived and planned the study. MB and SP wrote the original research code with MB contributing the majority of analysis and coding. All authors contributed to analyses and discussions. MB, SP and DP wrote the manuscript.

## References

1. Van Vreeswijk, C. & Sompolinsky, H. Chaos in neuronal networks with balanced excitatory and inhibitory activity. Science 274, 1724–1726 (1996). 10.1126/science.274.5293.1724

2. Haider, B., Duque, A., Hasenstaub, A.R. & McCormick, D.A. Neocortical network activity in vivo is generated through a dynamic balance of excitation and inhibition. The Journal of Neuroscience 26, 4535–4545 (2006). 10.1523/JNEUROSCI.5297-05.2006

3. Singer, W. & Effenberger, F. Oscillations in natural neuronal networks; an epiphenomenon or a fundamental computational mechanism? Human Arenas 8, 846–868 (2025). 10.1007/s42087-025-00478-x

4. Ermentrout, G. B. & Kleinfeld, D. Traveling electrical waves in cortex: insights from phase dynamics and speculation on a computational role. Neuron 29, 33–44 (2001). 10.1016/S0896-6273(01)00178-7

5. Muller, L., Busch, A. N., Davis, Z. W. & Reynolds, J. H. Neural traveling waves in cortex: Network mechanisms and potential roles in neural computation. Neuron 114, 3156–3174 (2026). 10.1016/j.neuron.2026.06.019

6. Plenz, D., Ribeiro, T. L., Miller, S. R., Kells, P. A., Vakili, A. & Capek, E. L. Self-organized criticality in the brain. Frontiers in Physics 9, 639389 (2021). 10.3389/fphy.2021.639389

7. Plenz, D. Neuronal avalanches and coherence potentials. The European Physical Journal Special Topics 205, 259–301 (2012). 10.1140/epjst/e2012-01575-5

8. Beggs, J.M. & Plenz, D. Neuronal avalanches in neocortical circuits. J. Neurosci. 23, 11167–11177 (2003). 10.1523/JNEUROSCI.23-35-11167.2003

9. Gardner, M. The fantastic combinations of John Conway’s new solitaire game “life”. Scientific American 223, 120–123 (1970). 10.1038/scientificamerican1070-120

10. Adamatzky, A. Game of Life on phyllosilicates: Gliders, oscillators and still life. Physics Letters A 377, 1597–1605 (2013). https://arxiv.org/abs/1306.0253

11. Eppstein, D. Searching for spaceships. In More Games of No Chance (ed. Nowakowski, R. J.), 433–453 (Cambridge University Press, 2002). https://library.slmath.org/books/Book42/files/eppstein.pdf

12. Johnston, N. The B36/S125 “2x2” life-like cellular automaton. Preprint at https://arxiv.org/abs/1203.1644v1 (2012).

13. Rennard, J.-P. Implementation of logical functions in the Game of Life. In Collision-Based Computing (ed. Adamatzky, A.), 491–512 (Springer, 2002). https://arxiv.org/abs/cs/0406009

14. Berlekamp, E. R., Conway, J. H. & Guy, R. K. Winning Ways for Your Mathematical Plays (Academic Press, 1982).

15. Lizier, J. T., Prokopenko, M. & Zomaya, A. Y. Local information transfer as a spatiotemporal filter for complex systems. Physical Review E 77, 026110 (2008). 10.1103/PhysRevE.77.026110

16. Crutchfield, J. P. & Mitchell, M. The evolution of emergent computation. Proc. Natl Acad. Sci. USA 92, 10742–10746 (1995). 10.1073/pnas.92.23.10742

17. Thiagarajan, T.C., Lebedev, M.A., Nicolelis, M.A. & Plenz, D. Coherence potentials: Loss-less, all-or-none network events in the cortex. PLoS Biology 8, e1000278 (2010). 10.1371/journal.pbio.1000278

18. Diesmann, M., Gewaltig, M.O. & Aertsen, A. Stable propagation of synchronous spiking in cortical neural networks. Nature 402, 529–533 (1999). 10.1038/990101

19. Aviel, Y., Horn, D. & Abeles, M. Synfire waves in small balanced networks. Neurocomputing 58-60, 123–127 (2004). 10.1016/s0925-2312(04)00028-1

20. Adamatzky, A. Localizations in cellular automata with mutualistic excitation rules. Chaos, Solitons & Fractals 40, 981–1003 (2009). 10.1016/j.chaos.2007.08.085

21. Peled, B. Y., Mishra, V. K. & Carmi, A. Y. Computing by nowhere increasing complexity. Preprint at https://arxiv.org/abs/1710.01654v1 (2017).

22. Martínez, G. J., Adamatzky, A. & McIntosh, H. V. Localization dynamics in a binary two-dimensional cellular automaton: The Diffusion Rule. Preprint at https://arxiv.org/abs/0908.0828v1 (2009).

23. Davis, Q. T. & Bongard, J. Selecting continuous life-like cellular automata for halting unpredictability: Evolving for abiogenesis. In Proceedings of the Genetic and Evolutionary Computation Conference Companion, 104–107 (2022). 10.1145/3520304.3529037

24. Kojima, H., Yevenko, I. & Ikegami, T. The glider equation for Asymptotic Lenia. Preprint at https://arxiv.org/abs/2508.04167v2 (2025).

25. Gong, P. & van Leeuwen, C. Distributed dynamical computation in neural circuits with propagating coherent activity patterns. PLoS Computational Biology 5, e1000611 (2009). 10.1371/journal.pcbi.1000611

26. Gong, P. & Robinson, P. A. Dynamic pattern formation and collisions in networks of excitable elements. Physical Review E 85, 055101(R) (2012). 10.1103/PhysRevE.85.055101

27. Chen, G. & Gong, P. Computing by modulating spontaneous cortical activity patterns as a mechanism of active visual processing. Nature Communications 10, 4915 (2019). 10.1038/s41467-019-12918-8

28. Baumgarten, L. & Bornholdt, S. Universal computation using localized limit-cycle attractors in neural networks. International Journal of Unconventional Computing 17, 239–258 (2022). https://www.oldcitypublishing.com/journals/ijuc-home/ijuc-issue-contents/ijuc-volume-17-number-4-2022/ijuc-17-4-p-239-258/

29. Maass, W., Natschläger, T. & Markram, H. Real-time computing without stable states: A new framework for neural computation based on perturbations. Neural Computation 14, 2531–2560 (2002). 10.1162/089976602760407955

30. Nichele, S. & Molund, A. Deep learning with cellular automaton-based reservoir computing. Complex Systems 26, 319–340 (2017). 10.25088/ComplexSystems.26.4.319

31. Legenstein, R. & Maass, W. Edge of chaos and prediction of computational performance for neural circuit models. Neural Networks 20, 323–334 (2007). 10.1016/j.neunet.2007.04.017

32. Tononi, G., Sporns, O. & Edelman, G. M. A measure for brain complexity: relating functional segregation and integration in the nervous system. Proc. Natl Acad. Sci. USA 91, 5033–5037 (1994). 10.1073/pnas.91.11.5033

33. Tononi, G., Sporns, O. & Edelman, G. M. A complexity measure for selective matching of signals by the brain. Proc. Natl Acad. Sci. USA 93, 3422–3427 (1996). 10.1073/pnas.93.8.3422

34. Mediano, P. A. M., Rosas, F. E., Luppi, A. I., Jensen, H. J., Seth, A. K., Barrett, A. B., Carhart-Harris, R. L. & Bor, D. Greater than the parts: a review of the information decomposition approach to causal emergence. Philosophical Transactions of the Royal Society A 380, 20210246 (2022). 10.1098/rsta.2021.0246

35. Gómez Soto, J. M. & Wuensche, A. Minimal glider-gun in a 2D cellular automaton. Preprint at https://arxiv.org/abs/1709.02655v1 (2017).

36. Gómez Soto, J. M. & Wuensche, A. X-Rule’s precursor is also logically universal. Preprint at https://arxiv.org/abs/1611.08829v1 (2016).

37. Gómez Soto, J. M. & Wuensche, A. The X-rule: Universal computation in a non-isotropic Life-like cellular automaton. Preprint at https://arxiv.org/abs/1504.01434v2 (2015).

38. Albantakis, L. et al. Integrated information theory (IIT) 4.0: Formulating the properties of phenomenal existence in physical terms. PLoS Computational Biology 19, e1011465 (2023). 10.1371/journal.pcbi.1011465

39. Tremblay, R., Lee, S. & Rudy, B. GABAergic interneurons in the neocortex: From cellular properties to circuits. Neuron 91, 260–292 (2016). 10.1016/j.neuron.2016.06.033

40. Wolfram, S. Statistical mechanics of cellular automata. Reviews of Modern Physics 55, 601–644 (1983). 10.1103/RevModPhys.55.601

41. Greenberg, J.M. & Hastings, S.P. Spatial Patterns for Discrete Models of Diffusion in Excitable Media. SIAM Journal on Applied Mathematics 34, 515–523 (1978). 10.1137/0134040

42. Pfeffer, C.K., Xue, M., He, M., Huang, Z.J. & Scanziani, M. Inhibition of inhibition in visual cortex: the logic of connections between molecularly distinct interneurons. Nature Neuroscience 16, 1068–1076 (2013). 10.1038/nn.3446

43. Jiang, X. et al. Principles of connectivity among morphologically defined cell types in adult neocortex. Science 350, aac9462 (2015). 10.1126/science.aac9462

44. Rybarsch, M. & Bornholdt, S. Avalanches in self-organized critical neural networks: A minimal model for the neural SOC universality class. PLoS One 9, e93090 (2014). 10.1371/journal.pone.0093090

