## Supplemental Material for "Glider compositionality and information capacity in an E/I-balanced neuronal automaton"

#### Supplementary Note 1. Reduced dynamics and propagation boundaries of the G-glider

##### Reduction to an amplitude map

The G-glider repeats its spatial pattern after four time steps, with a reflection after two. Only a subset of active cells drives the next propagation step (main Fig. 3a–c). In the minimal state, one excitatory component supplies the cells needed for the next G-shaped state. Two excitatory inputs and one inhibitory input then determine the next minimal state. If the spatial and refractory pattern is maintained and no other activity interferes, these interactions give

$$e_1^3 = \sigma\left(\frac{S_{EE}}{9}(e_1^2 + e_2^2) - \frac{S_{IE}}{5} i_1^2\right)$$

$$e_1^2 = \sigma\left(\frac{S_{EE}}{9} e_1^1\right) = e_2^2$$

$$i_1^2 = \sigma\left(\frac{S_{EI}}{20} e_1^1\right)$$

Substituting the component responses yields

$$\Rightarrow e_1^3 = \sigma\left(\frac{2S_{EE}}{9}\sigma\left(\frac{S_{EE}}{9}e_1^1\right) - \frac{S_{IE}}{5}\sigma\left(\frac{S_{EI}}{20}e_1^1\right)\right)$$

For the principal parameter plane, let $E = S_{EE} = S_{EI}$ and $I = S_{IE}$. These symbols denote coupling strengths in this derivation, rather than the cellular activity fields used in the Methods. The component activity x then evolves through the map

$$f(x) = \sigma\left(\frac{2E}{9}\sigma\left(\frac{E}{9}x\right) - \frac{I}{5}\sigma\left(\frac{E}{20}x\right)\right)$$

where

$$\sigma(x) = \tanh\left(\frac{x}{2}\right)\ for\ x \geq 0, \qquad 0\ \ otherwise$$

One iteration advances the component amplitude by two network time steps. At a fixed point, the component amplitude repeats across successive half-cycles. Inhibitory–inhibitory coupling does not enter this reduction for the assumed isolated G-glider cycle; this does not imply that it is irrelevant to the surrounding network or glider collisions.

##### Fixed points and the active branch

Write the input to the outer transfer function as g(x), so f(x) = σ(g(x)). Define $P = 2E^2/81$ and $Q = IE/100$. At differentiable points on the active branch, the chain rule gives

$$f'(x) = \sigma'(g(x))[P\sigma'\left(\frac{Ex}{9}\right) - Q\sigma'\left(\frac{Ex}{20}\right)]$$

For positive arguments, $\sigma'(z) = \frac{1}{2}\,\mathrm{sech}^2(z/2)$. The rectified function has no two-sided derivative at zero; we use right derivatives on the nonnegative activity domain. For $E > 0$ and $I < 200E/81$,

$$f'(0+) = \frac{1}{4}\left(\frac{2E^2}{81} - \frac{IE}{100}\right)$$

For $I \geq 200E/81$, the map is identically zero for nonnegative activity. Extending the active-branch expression into this region yields negative values that are not derivatives of the rectified map.

The derivative bound follows from $\sigma'(Ex/9) \leq \sigma'(Ex/20)$ for $x \geq 0$. With $A = \sigma'(Ex/9)$, $B = \sigma'(Ex/20)$, the inner derivative is $a_2 = PA - QB$ and satisfies

$$a_2 = PA - QB \leq (P - Q)B$$

because $P(B-A) \geq 0$. In the domain $P-Q > 0$, $B$ is bounded above by $\sigma'(0+)$. If $a_2 \geq 0$, the outer derivative is also bounded above by $\sigma'(0+)$, giving $f'(x) \leq f'(0+)$. If $a_2 < 0$, $f'(x) \leq 0$, which satisfies the same upper bound. The inactive branch has zero derivative; boundary points are evaluated using one-sided derivatives.

For $E > 0$, the map is continuous on the activity interval, $f(0) = 0$ and $f(1) < 1$. A right derivative greater than one therefore guarantees a positive fixed point. Solving this condition gives

$$I < \frac{200E}{81} - \frac{400}{E}$$

This is a condition for existence of a nonzero fixed point of the reduced map. Local stability additionally requires $|f'(x^*)| < 1$. Higher-period amplitude orbits require their own stability test. The fixed-point condition alone does not establish preservation of the assumed spatial pattern.

We can also solve for the condition when $f(x)$ is identically zero everywhere by noting that $f(x)>=0$ for all $x$ and $f'(x)$ is maximized at $x=0+$. Thus, for $f'(0+)<=0$ we know $f(x)$ is identically zero. Solving this condition gives

$$I \geq \frac{200E}{81}$$

#### Activation outside the glider pattern

Cells outside the assumed G-glider pattern can become active and disrupt propagation (main Fig. 3c). For the checkpoint denoted $x_1$, the source reduction gives

$$x_1 = \sigma\left(\frac{2E}{9}\sigma\left(\frac{E}{9}e_1^2\right) - \frac{2I}{5}\sigma\left(\frac{E}{20}e_1^2\right)\right)$$

For a positive component amplitude $z$, the sign of its incoming drive can be evaluated as

$$\frac{2E}{9}\sigma\left(\frac{Ez}{9}\right) - \frac{2I}{5}\sigma\left(\frac{Ez}{20}\right) = \frac{2E}{9}\tanh\left(\frac{Ez}{18}\right) - \frac{2I}{5}\tanh\left(\frac{Ez}{40}\right)$$

The transfer-function arguments are halved only when $\sigma$ is replaced by tanh. Since $\tanh(Ez/18) > \tanh(Ez/40)$ for $E,z > 0$, $E > 9I/5$ is a sufficient condition for positive activation at this checkpoint. Comparing the derivatives of the two terms gives a sufficient condition for no activation, $I > 100E/81$. These bounds concern this checkpoint; the complete spatial constraints also depend on refractoriness and the other candidate sites.

Analytical bounds, numerical iteration, and observed network behavior contribute different boundaries to main Fig. 3. They do not constitute a single analytical classification of every dynamical region. The period-doubling structure in Supplementary Fig. 4 concerns the reduced amplitude map, whereas the network catalogue concerns spatially recurring patterns.

#### Transfer gain

For $\sigma_a(z) = \sigma_1(az)$, the reduced dynamics satisfy $f_a(x;E,I) = f_1(x;aE,aI)$. Changing gain therefore rescales the coupling values for corresponding dynamics (Supplementary Fig. 7). This is distinct from changing transfer-function shape (Supplementary Fig. 8) and does not by itself establish a new threshold for mixed-glider composition.

#### 64 Supplementary Figure 1

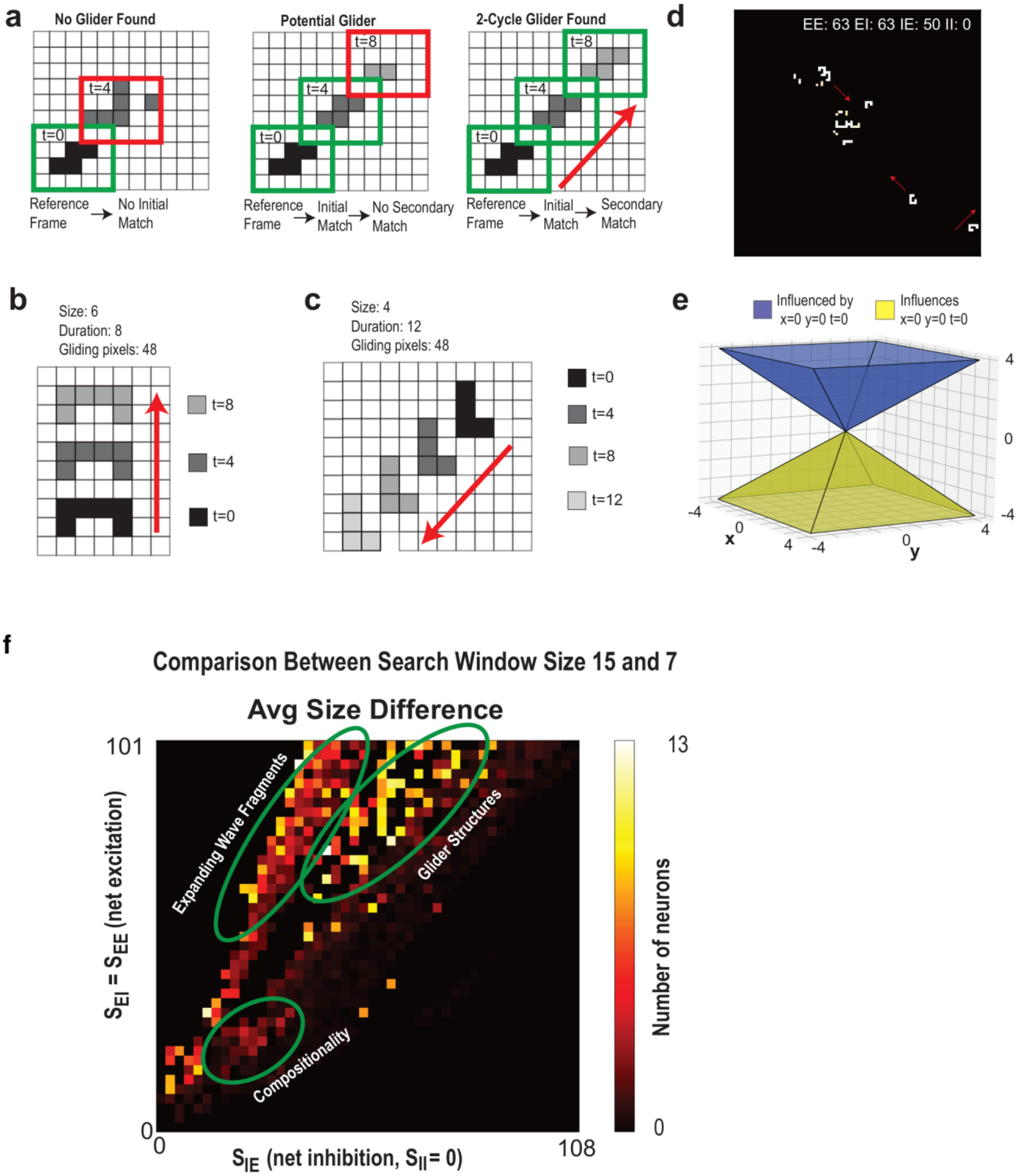

Supplementary Figure 1 | Detection of spatially recurring activity patterns and dependence on search-window size. (a) A candidate must recur after τ and 2τ at the same displacement per interval; failure at either comparison rejects it. (b,c) U- and L-glider examples with period, size and observed duration. (d) Excitatory activity and detected trajectories at the displayed coupling tuple. (e) Search geometry associated with the imposed velocity constraint. The implementation binarizes activity above zero, searches windows of 3–10 cells in each dimension and recurrence intervals of 1–4 time steps, requires a silent window border and rejects zero displacement. Patterns exceeding these spatial or temporal limits are excluded from the catalogue. (f) Difference in mean detected pattern size between maximum search-window sizes of 15 and 7 in independent simulations across the excitation–inhibition parameter plane at zero inhibitory–inhibitory coupling. Color indicates the difference in mean number of neurons excluding cases with no glider detections and points where the difference of mean size was negative. Marked regions identify compositionality, glider structures and expanding wave fragments. The larger window detects patterns with more neurons in several regions, including the compositionality region.

#### Supplementary Figure 2

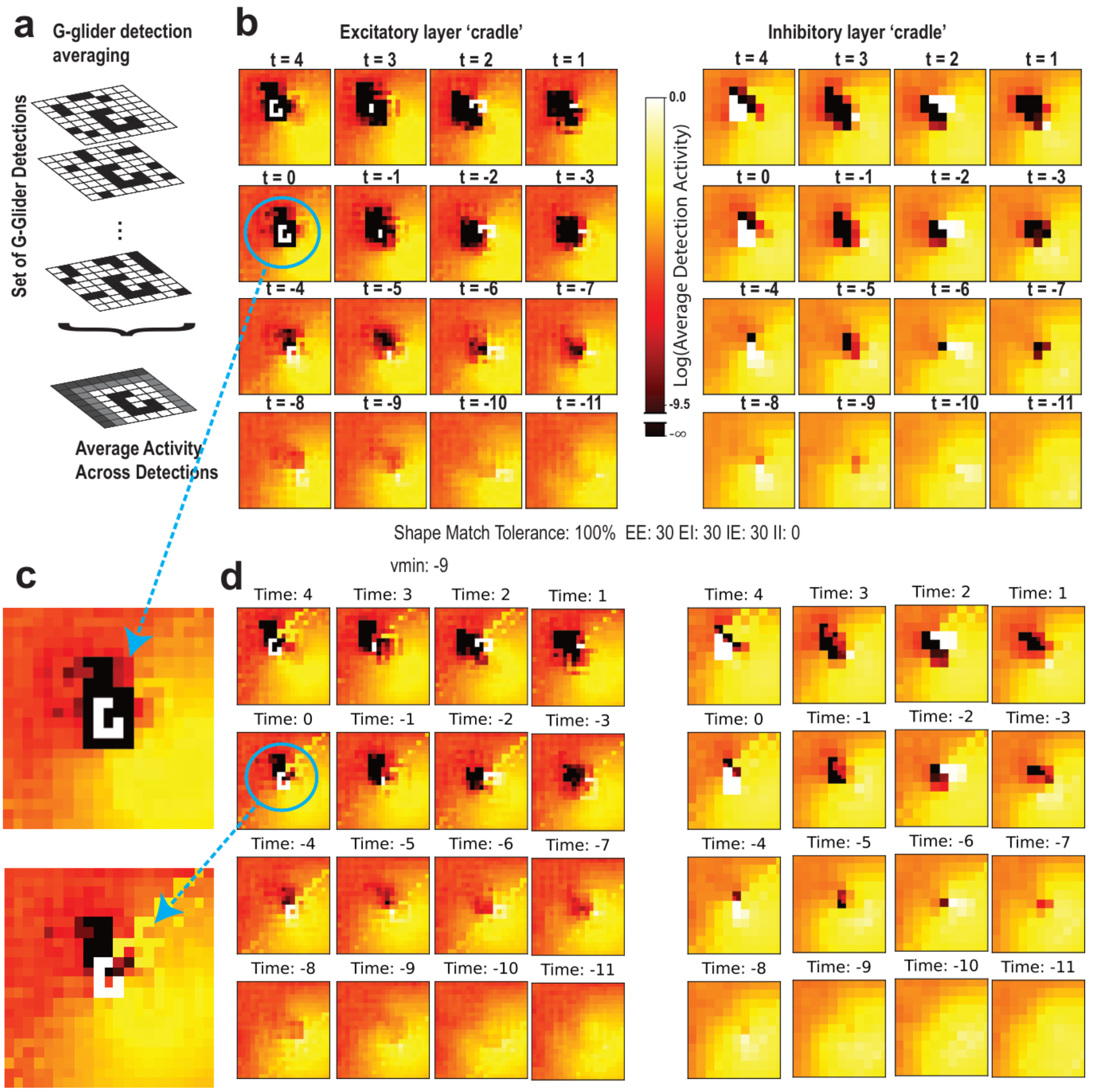


Supplementary Figure 2 | Reduced activity precedes G-glider detection. (a) Alignment of detected G-gliders and their surrounding activity by rotation and reflection. (b) Mean excitatory and inhibitory activity before and after detection, at $S_{EE} = S_{EI} = S_{IE} = 30$ and $S_{II} = 0$. The analysis includes 12,866 detections across 100 simulations of 3,000 time steps. Color denotes log10 mean activity; black indicates zero. Reduced activity ahead of emergence becomes apparent seven to eight steps before detection. (c) Enlarged emergence patterns. (d) A similar quiescent region precedes detection when adjacent active cells are allowed and the shape-matching threshold is relaxed to 90% rather than 100%. This comparison addresses a detector constraint but does not establish causal preparation of a route.

### Supplementary Figure 3

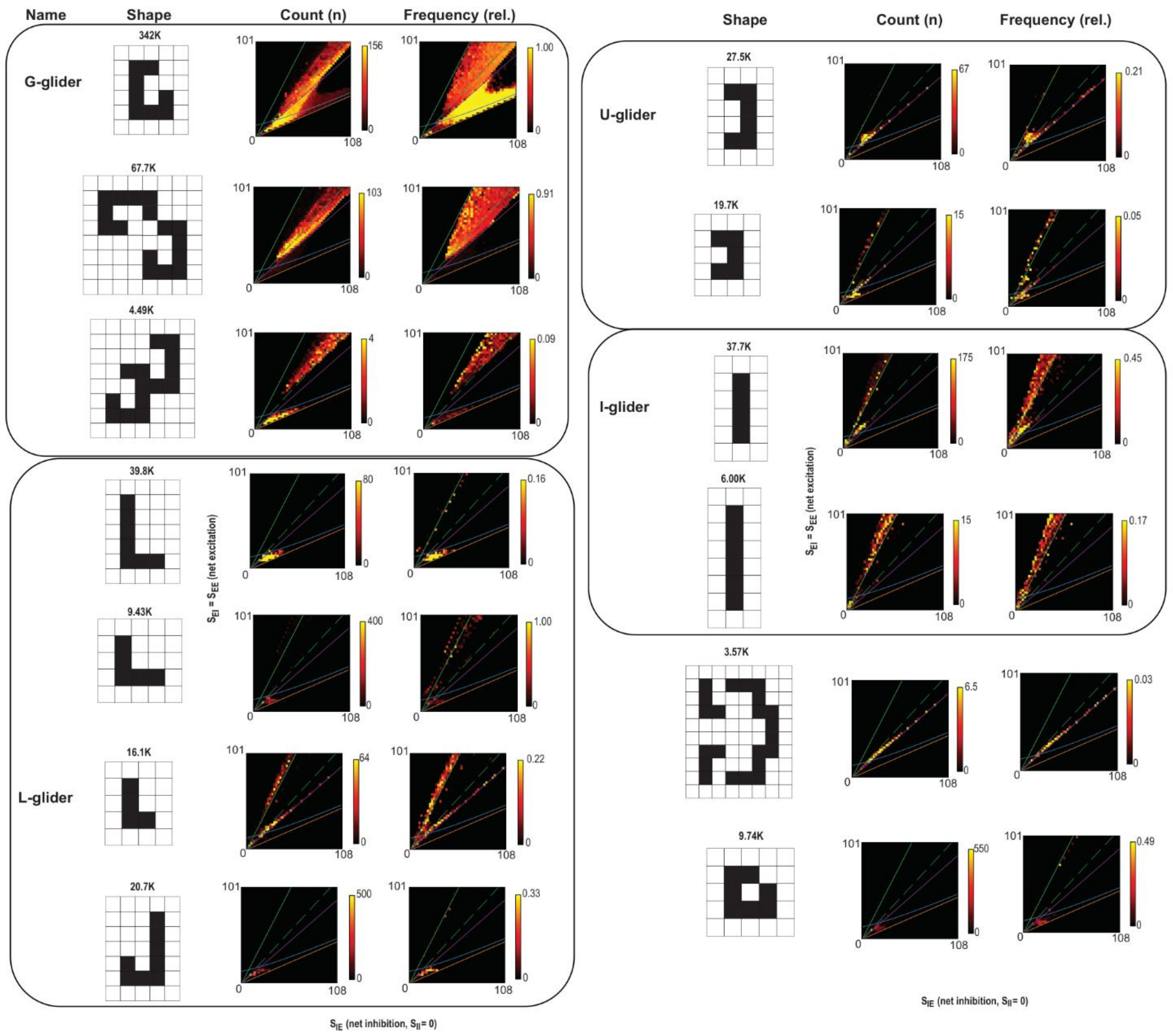


Supplementary Figure 3 | Pattern-specific abundance and dominance across parameter space. Shapes, detection counts, and relative frequencies are shown for thirteen frequently detected configurations with $S_{EE} = S_{EI}$ and $S_{II} = 0$. Relative frequency is normalized by total detections at each parameter point. The G-glider predominates along much of the equal-coupling region, while other configurations occupy neighboring domains. These relative frequencies depend on the finite search windows and recurrence intervals and do not establish dominance over larger patterns outside the search range.

### Supplementary Figure 4

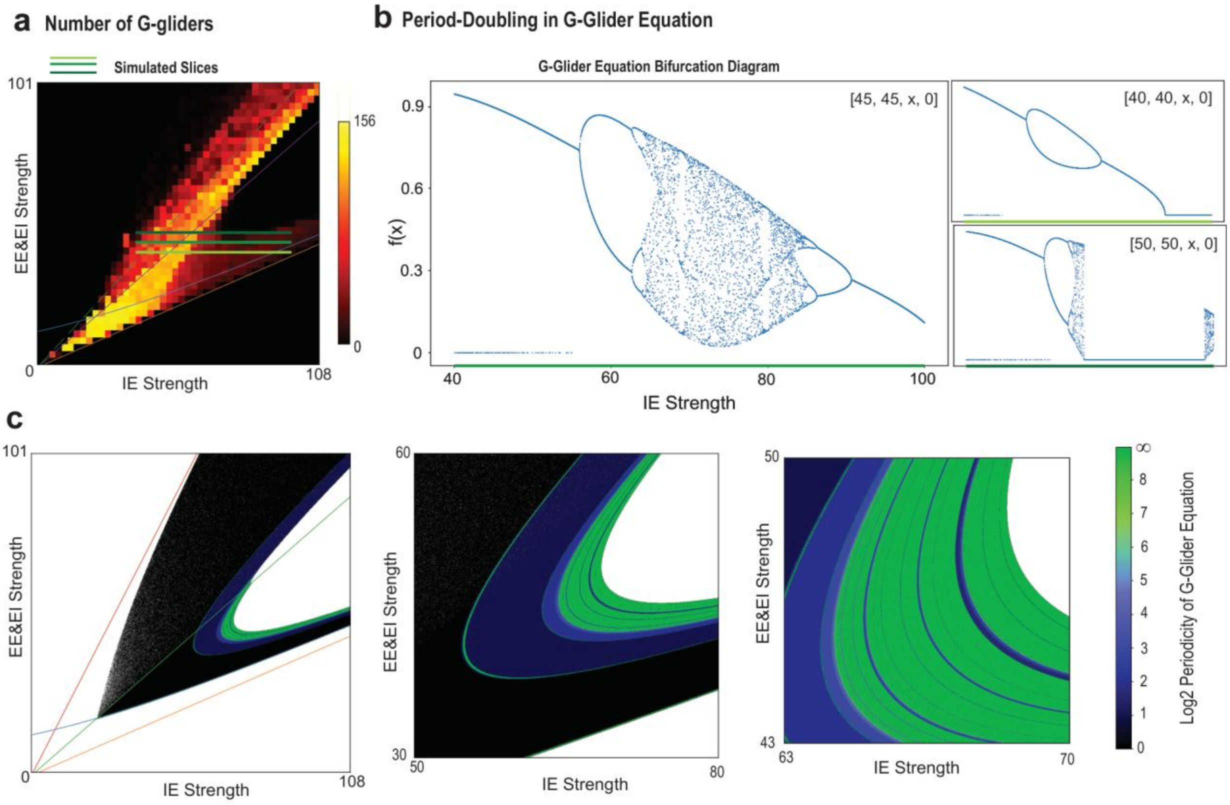


Supplementary Figure 4 | Bifurcations in the reduced G-glider amplitude map. (a) Parameter slices overlaid on simulated G-glider counts. (b) Bifurcation diagrams along those slices. (c) Periodicity of the amplitude map, including enlarged period-doubling bands. White points denote unstable iterates. Periodicity of the amplitude map does not establish stability of the full spatial glider pattern.

### Supplementary Figure 5

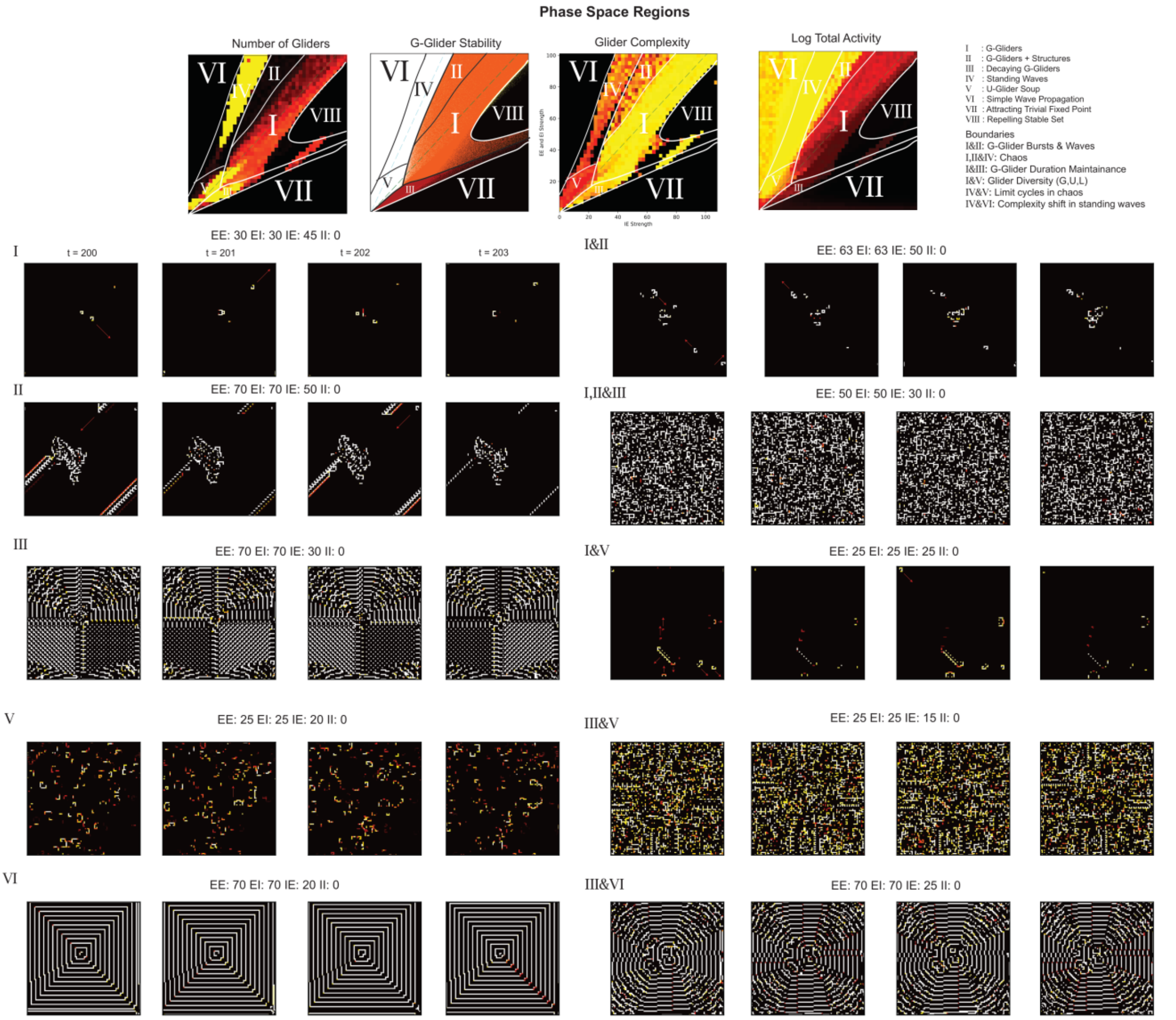


Supplementary Figure 5 | Dynamical regimes and boundary patterns. Maps compare detected gliders, reduced-map stability, pattern size and total activity. Four-step snapshot sequences illustrate nonquiescent regions and selected boundaries, with coupling values printed beside each sequence. Region labels follow main Fig. 3: III denotes decaying G-gliders and IV standing waves. Boundary examples include growing G-glider structures, mixed G/U activity and wave transitions.

104 Supplementary Figure 6

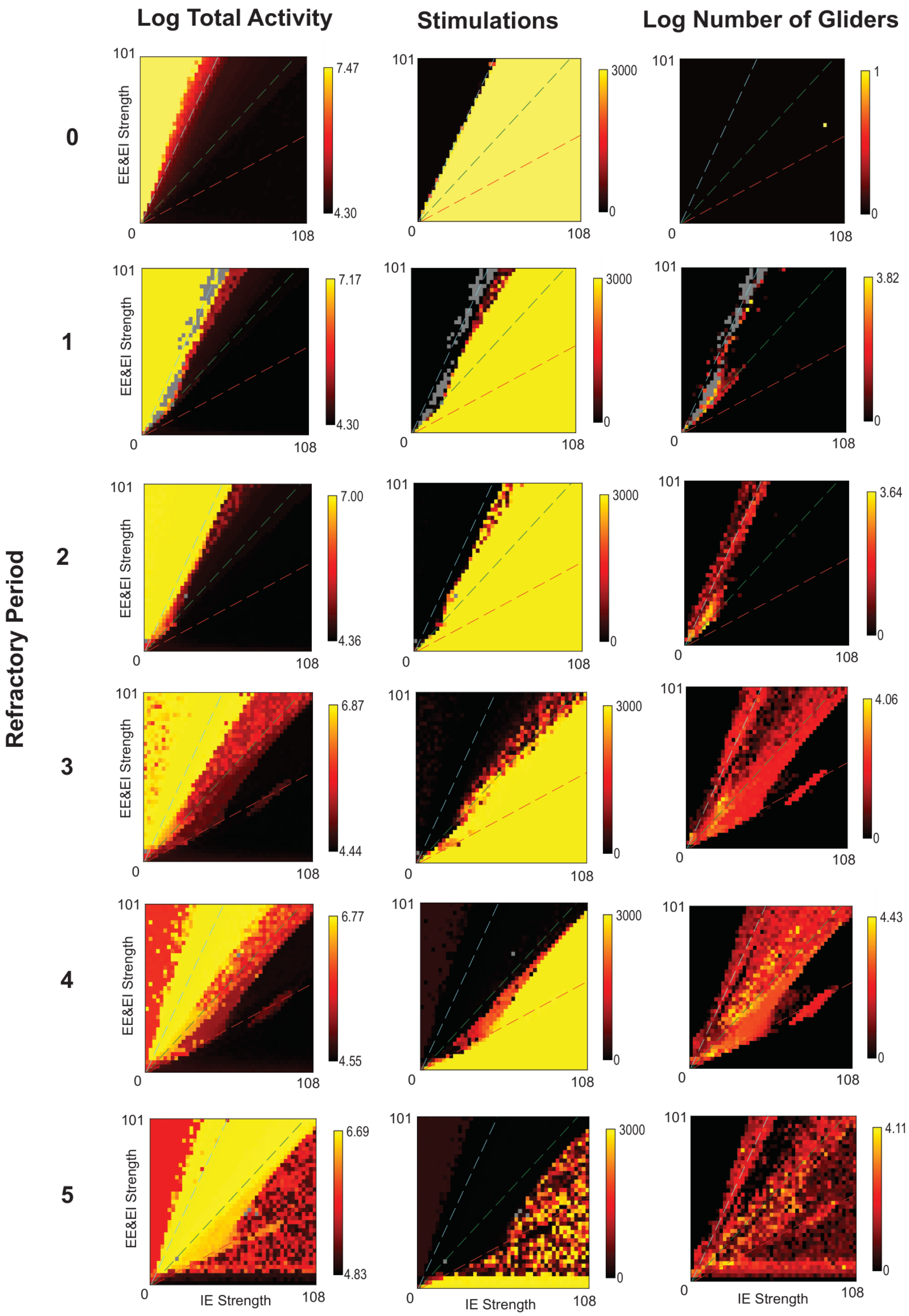

Supplementary Figure 6 | Refractoriness changes the domains of propagating activity. Rows show refractory periods of zero through five time steps. Columns show total activity, stimulation count, and detected-pattern count across $S_{EE} = S_{EI}$ versus $S_{IE}$ at $S_{II} = 0$, using the indicated transforms. No gliders were identified without refractoriness in this exploration. For periods one and two, some detections in the transition zone were classified as standing-wave fragments; isolated gliders occurred at lower coupling. Refractory periods of at least three time steps support a broader regime of rich glider activity within the explored range. At period five, the observed repertoire shifts toward larger structures and wave-associated patterns.

### Supplementary Figure 7

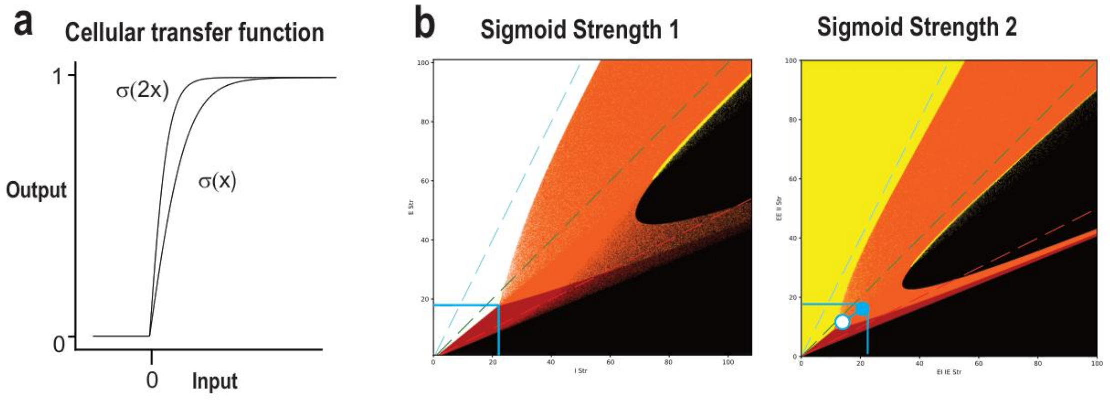


Supplementary Figure 7 | Transfer gain rescales the coupling strengths of reduced-map dynamics. (a) Transfer functions at the two displayed gains. (b) Corresponding reduced-map stability diagrams. With $\sigma_a(z) = \sigma_1(az)$, the map satisfies $f_a(x;E,I) = f_1(x;aE,aI)$, explaining the rescaling of corresponding coupling regions. This comparison tests gain–coupling equivalence rather than a separate compositionality threshold.

### Supplementary Figure 8

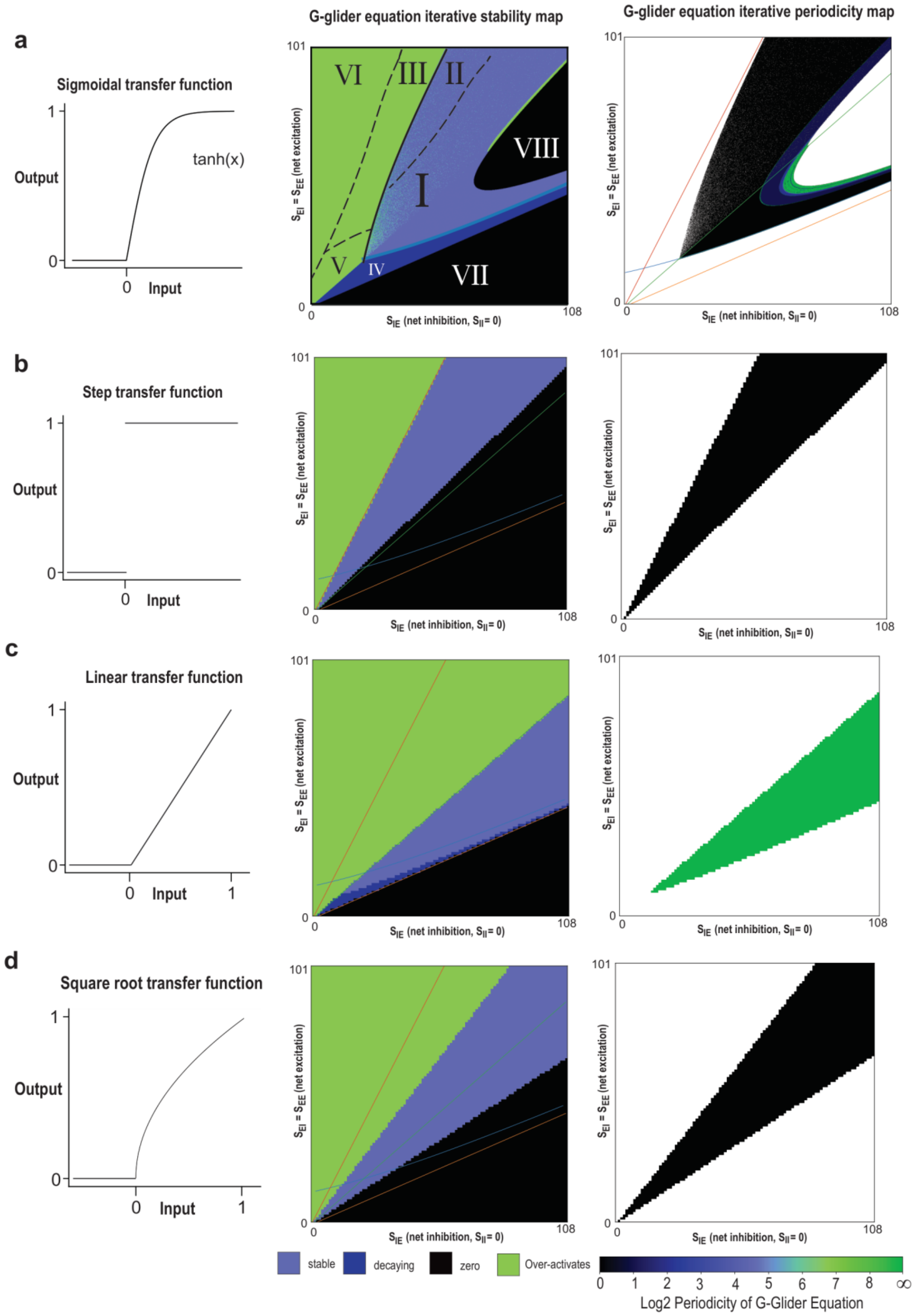


Supplementary Figure 8 | Transfer-function shape changes reduced G-glider dynamics. Each row shows a transfer function, its iterative-dynamics map and its periodicity map: (a) sigmoid, (b) binary step, (c) linear and (d) square root. The binary step and square-root functions support fixed-point dynamics, whereas the tested linear variant produces unbounded amplification of active solutions and does not sustain bounded G-glider activity.

### Supplementary Figure 9

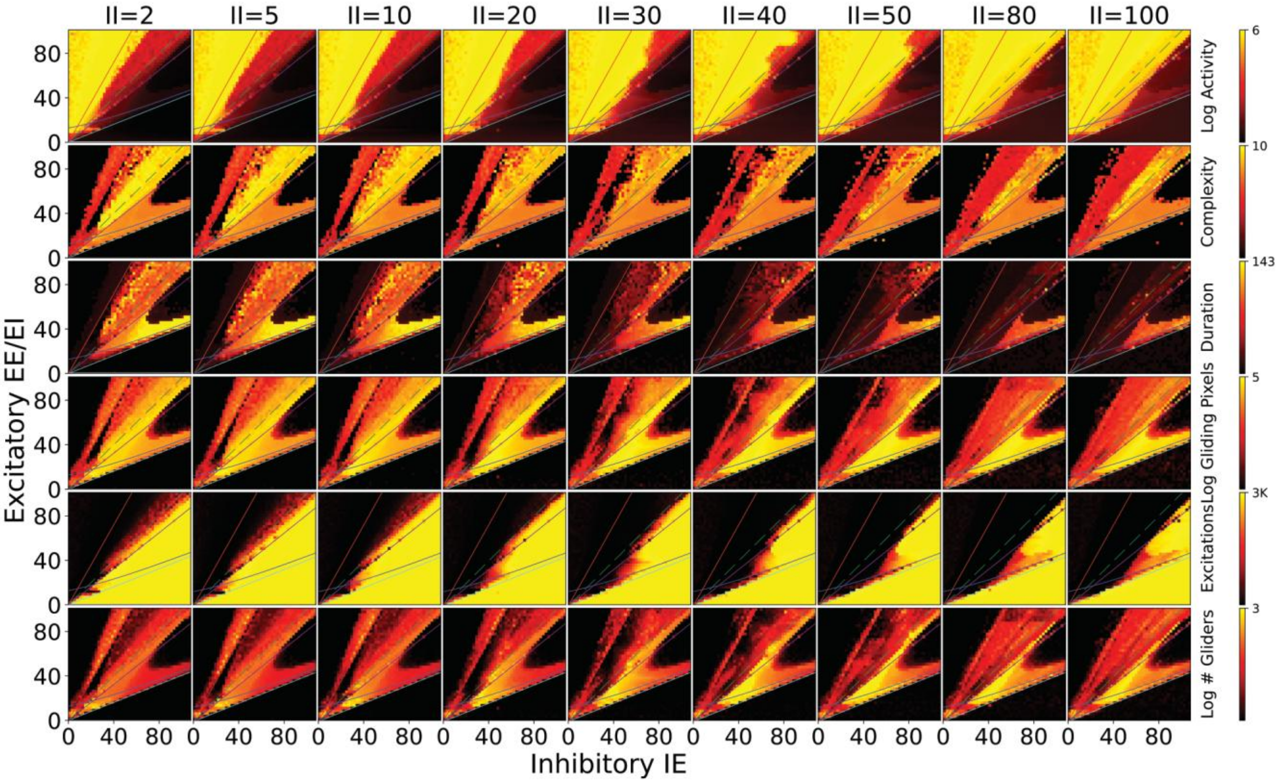


Supplementary Figure 9 | Network activity and detected patterns across inhibitory–inhibitory coupling. Columns show $S_{II}$ = 2, 5, 10, 20, 30, 40, 50, 80, and 100. Rows show the displayed measures of activity, pattern size, duration, cumulative gliding activity, stimulation, and detected-pattern count across $S_{EE} = S_{EI}$ versus $S_{IE}$. The broad regime organization persists, with shifts in the domains of waves and detected propagating patterns.

### Supplementary Figure 10

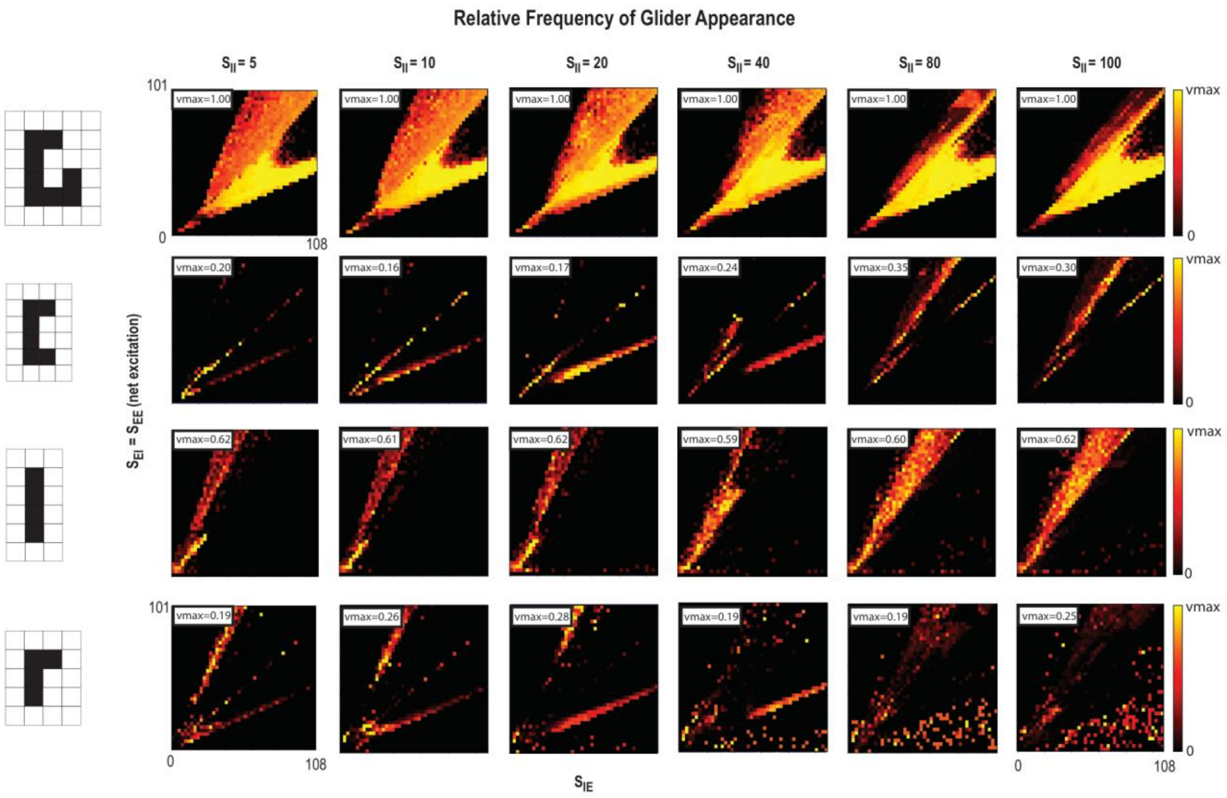


Supplementary Figure 10 | Pattern-specific dominance across inhibitory–inhibitory coupling. Rows show the G-, U-, I-, and L-shaped patterns; columns show $S_{II}$ = 5, 10, 20, 40, 80, and 100. Heatmaps report relative detection frequency across $S_{EE}$ = $S_{EI}$ versus $S_{IE}$. G- and I-pattern domains vary less than U- and L-pattern domains across the displayed conditions.

#### Supplementary Figure 11

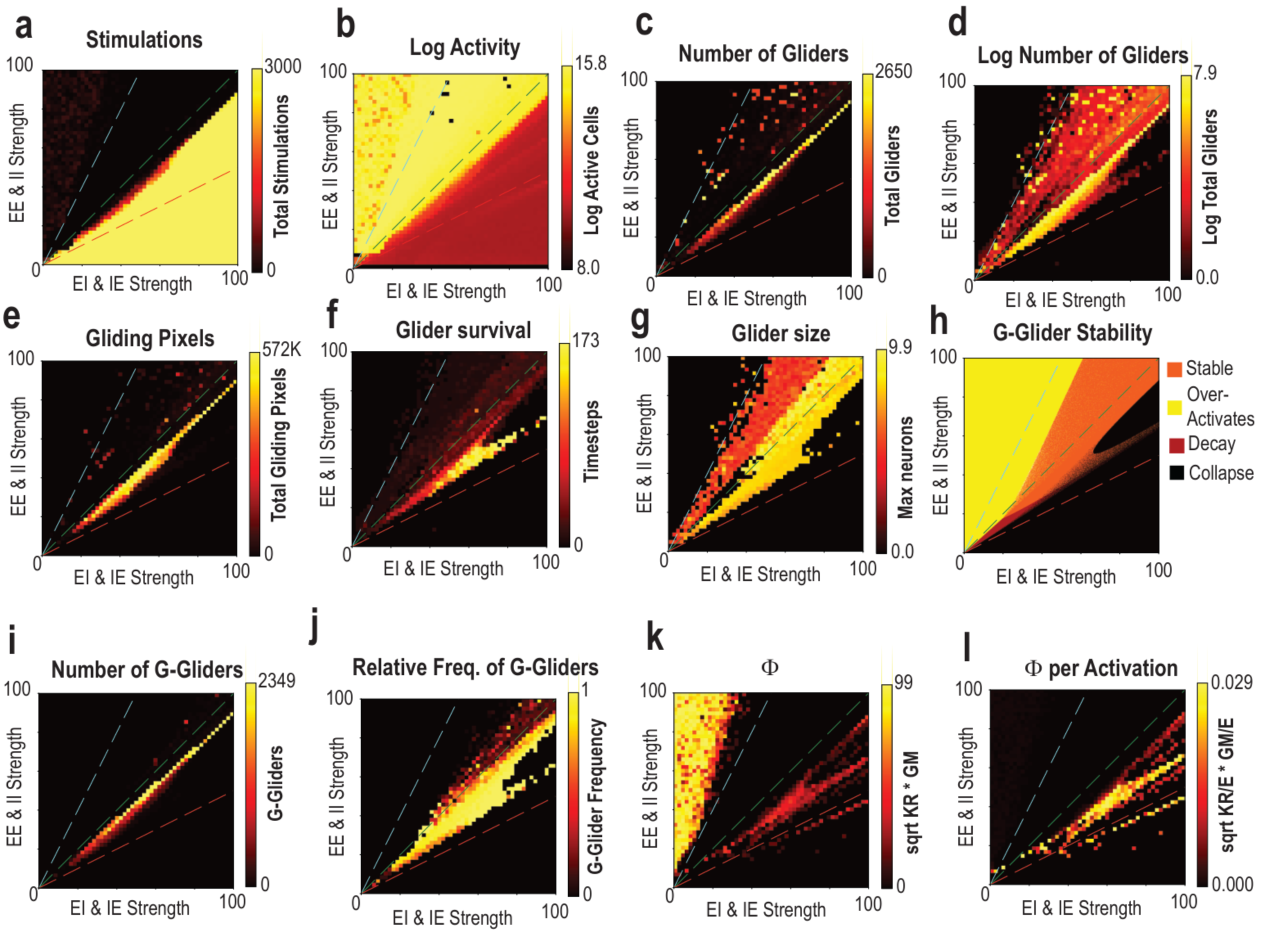


Supplementary Figure 11 | An alternative coupling plane resolves activity and glider domains. The vertical axis ties $S_{EE}$ to $S_{II}$ and the horizontal axis ties $S_{EI}$ to $S_{IE}$. (a,b) External stimulation and total activity. (c,d) Detected-pattern counts on linear and logarithmic scales. (e–g) Cumulative gliding activity, survival, and size. (h) Reduced G-glider stability. (i,j) G-glider count and relative frequency. (k,l) Rank-based capacity and activity-normalized capacity.

### Supplementary Figure 12

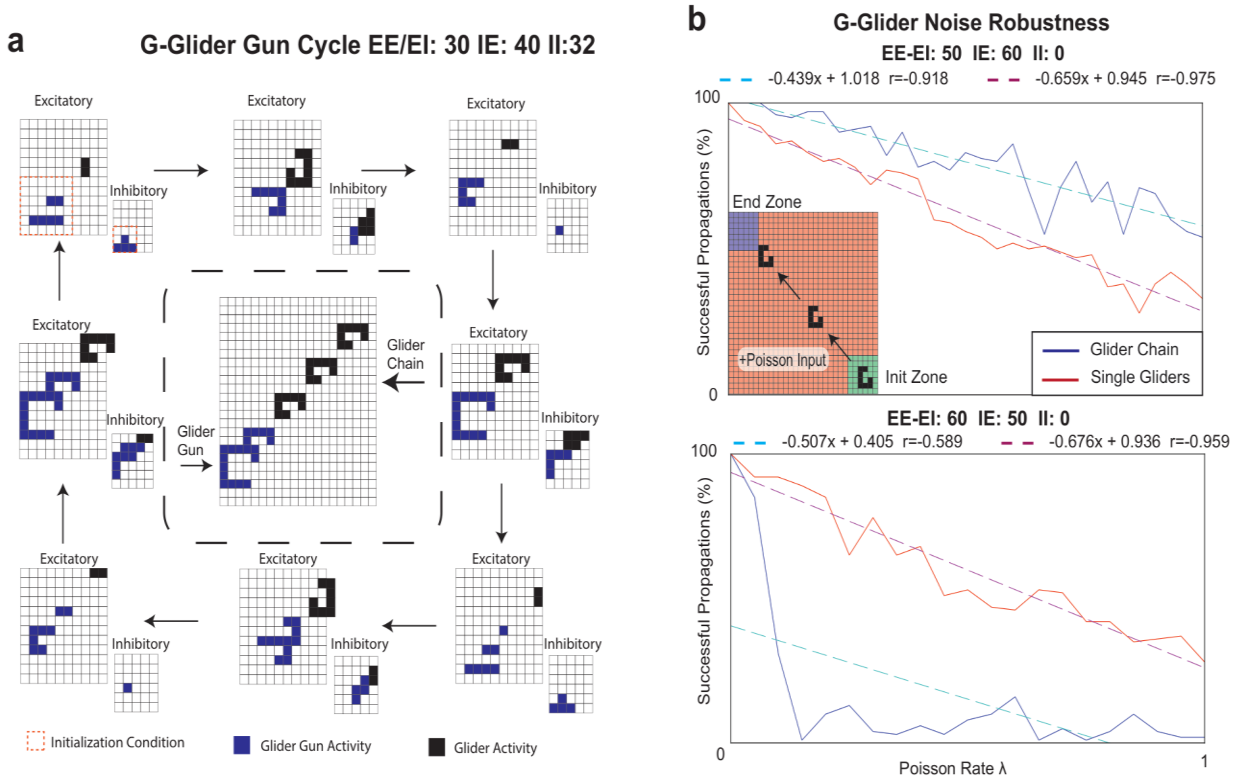


Supplementary Figure 12 | G-glider generation and transmission through noise. (a) Glider-source cycle at $S_{EE} = S_{EI} = 30$, $S_{IE} = 40$, $S_{II} = 32$, including initialization and emitted activity. (b) Transmission success for isolated gliders and chains at $(S_{EE},S_{EI},S_{IE},S_{II}) = (50,50,60,0)$, upper plot, and (60,60,50,0), lower plot. Chains are more robust in the first condition; isolated gliders are more robust in the second. λ is the mean number of attempted noise activations per network time step. Success requires shape-matched arrival at the expected time. For both isolated gliders and chains, success is scored per launched glider. The reported sampling comprised 100 independent single-glider trials and one chain of 100 gliders; gliders within the chain are not independent replicates. Curves and fits are distinguished; fitted intercepts are not observed probabilities.

### Supplementary Figure 13

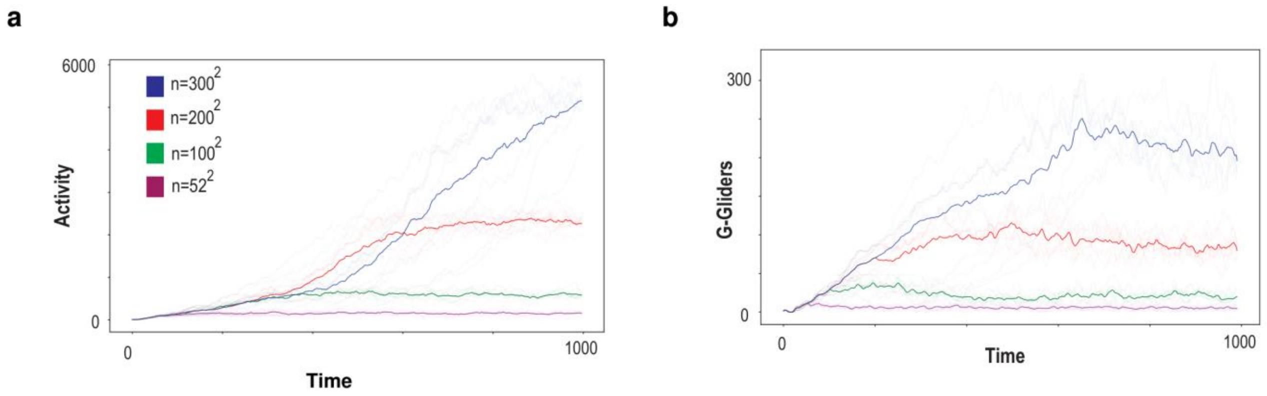


Supplementary Figure 13 | Sustained activity and glider counts depend on network size. (a) Activity and (b) G-glider-count trajectories at $(S_{EE},S_{EI},S_{IE},S_{II}) = (45,50,50,45)$. The displayed color key identifies excitatory grids of 52×52, 100×100, 200×200 and 300×300 cells. The tested 24×24 condition became quiescent and is not plotted. Thin traces represent individual collision arrangements; emphasized curves show their means. The size-analysis plotter uses four-step smoothing and selects sustained arrangements.

#### Supplementary Figure 14

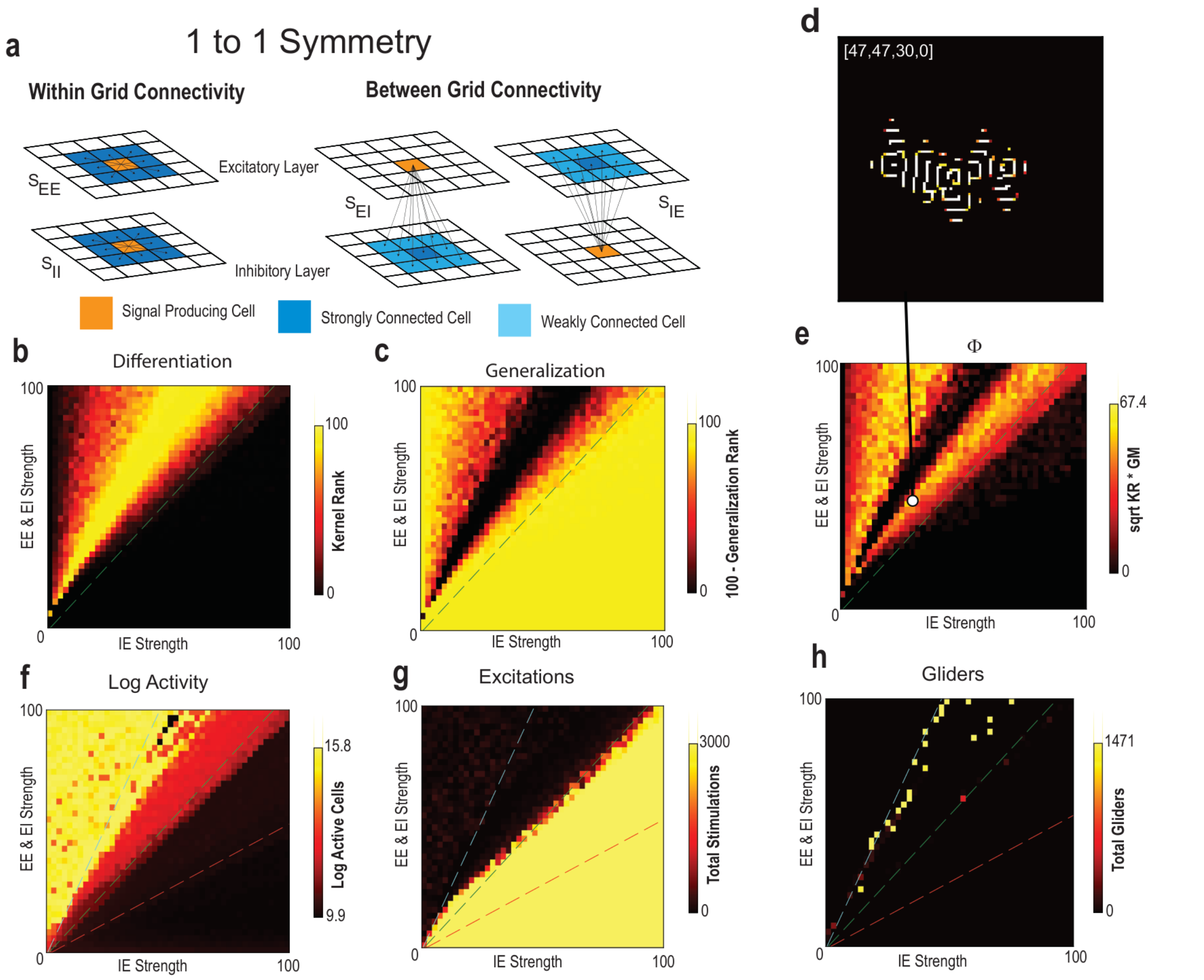


Supplementary Figure 14 | A symmetric 1:1 network supports capacity with wave-associated detector matches. (a) Symmetric within- and between-population connectivity. (b,c) Kernel rank and generalization measure. (d) Activity snapshot at the marked high-capacity setting (47,47,30,0). (e) Global Φ. (f,g) Total activity and external stimulation. (h) Detected-pattern counts; the source inspection identified standing-wave fragments rather than verified isolated gliders. The parameter plane uses $S_{EE}$ = $S_{EI}$ against $S_{IE}$, with $S_{II}$ = 0.

### Supplementary Figure 15

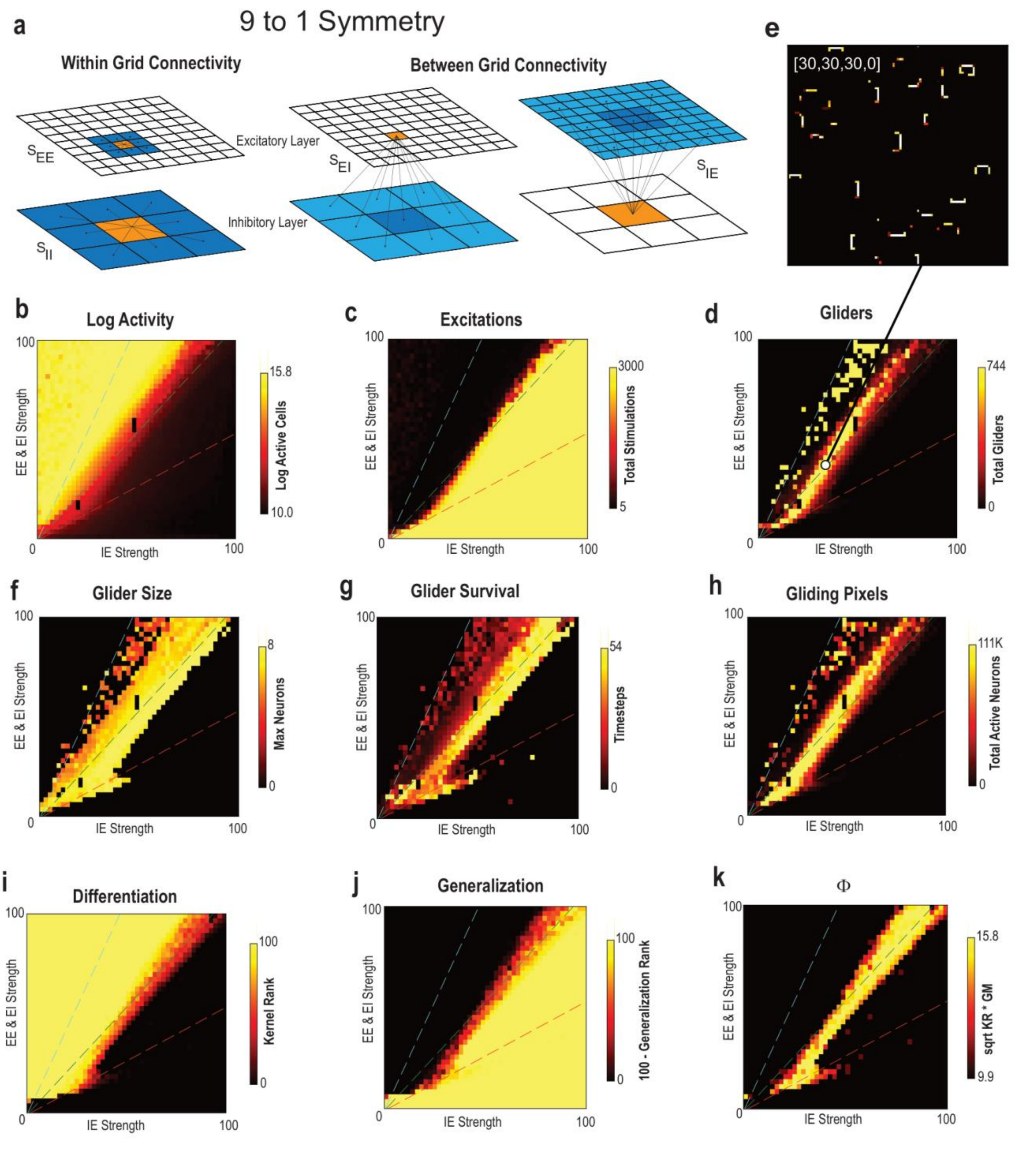


Supplementary Figure 15 | A symmetric 9:1 architecture also generates gliders. (a) Connectivity for the 9:1 population ratio: 99×99 excitatory and 33×33 inhibitory cells. (b,c) Activity and external stimulation. (d) Detected-glider count. (e) Activity snapshot at ($S_{EE}$,$S_{EI}$,$S_{IE}$,$S_{II}$) = (30,30,30,0). (f–h) Mean glider size, survival and cumulative gliding activity. (i–k) Kernel rank, generalization measure and Φ. The explored plane uses $S_{EE}$ = $S_{EI}$ against $S_{IE}$, with SII = 0. This control shows that the specific local asymmetry of the 4:1 model is not necessary for all glider emergence.
